# Human RIG-I deficiency confers susceptibility to Kaposi Sarcoma via loss of latency control

**DOI:** 10.1101/2025.09.19.676307

**Authors:** Lucie Roussel, Stéphane Bernier, Mélanie Langelier, Yichun Sun, Irem Ulku, Benjamin Mak, Yongbiao Li, Anna Perez, Alexi Wloski, Isabelle Angers, Lily-Rose Vinh, Annie Beauchamp, Sarah Boissel, A. Kevin Watters, Simon Rousseau, Jean-Pierre Routy, Virginie Calderon, Carolina Arias, Donald C. Vinh

**Affiliations:** Centre of Reference for Genetic Research in Infection and Immunity, Research Institute - McGill University Health Centre; Montreal, Quebec, Canada; Molecular biology and functional genomics Core, Montreal Clinical Research Institute (IRCM); Montreal, Quebec, Canada; Department of Pathology, McGill University Health Centre, Montreal, Quebec, Canada; Meakins-Christie Laboratories, Research Institute of McGill University Health Centre; Montreal, Quebec, Canada; Division of Hematology and Chronic Viral Illness Service, McGill University Health Centre: Glen Site Research Institute; Montreal, Quebec, Canada; Department of Molecular, Cellular, and Developmental Biology, University of California, Santa Barbara; Center for Stem Cell Biology and Engineering, University of California; Santa Barbara, United States of America; Department of Medicine (Division of Infectious Diseases), McGill University Health Centre; Department of OptiLab (Division of Medical Microbiology, Division of Molecular Genetics-Immunology), McGill University Health Centre; Department of Human Genetics, McGill University

**Keywords:** RIG-I, DDX58, Kaposi’s sarcoma-associated herpesvirus (KSHV), viral latency, interferon signaling, type I interferons, ISGs, viral oncogenesis, DNA virus sensing, innate immunity, IFN-β, IFN-ω

## Abstract

Kaposi sarcoma (KS), caused by the DNA-virus Kaposi’s sarcoma-associated herpesvirus (KSHV), occurs during T cell immunosuppression (HIV, transplant) or sporadically in some ‘immunocompetent’ and aging individuals (endemic, classic KS respectively). In absence of known T cell immunosuppression KS pathogenesis remains enigmatic. KS therapy with topical or oral retinoid medication, or recombinant alpha interferon, can induce remission and suggests the involvement of two signalling pathways. Retinoic acid-inducible gene-I (RIG-I) encoded by *DDX58* is canonically a sensor of RNA-viruses, its function in human immunity against DNA-viruses remains poorly defined. We report a patient with classic KS, carrying a homozygous nonsense (p.Q393*) mutation in *DDX58*, abolishing RIG-I expression and specifically impairing RIG-I agonist responses. In isogenic cell models, loss of RIG-I compromised responses during both KSHV primary infection and viral reactivation, diminishing induction of type I interferons and interferon-stimulated genes, skewing to a persistent latent viral gene program, and dysregulating cellular pro-oncogenic pathways by transcriptomic and proteomic profiling. This work defines the first innate immunodeficiency underlying classical KS, revealing RIG-I’s role in KSHV immunopathogenesis and expanding its function in human antiviral immunity beyond RNA-viruses, while identifying promising therapeutic targets.

**Significance statement:** RIG-I deficiency causes classic KS by failing to control KSHV infection and reactivation, expanding its role beyond RNA-viruses.

## INTRODUCTION

Kaposi’s sarcoma (KS), a vascular neoplasia originating from endothelial cells, is caused by Kaposi’s sarcoma associated herpesvirus (KSHV), also known as human herpesvirus 8 (HHV-8), a double-stranded DNA virus belonging to the gamma *Herpesviridae* family. Seroprevalence as surrogate marker of infection varies according to global geography, with highest rates in Central and Eastern Africa (>50%), intermediate rates in Western Africa, South America, Mediterranean basin, and Eastern Europe (10-30%), and lowest rates in Western Europe, North America, and Eastern Asia (<10%). Within these regions, geographical and socio-demographic factors as well as sexual behaviors may further contribute to heterogeneity in infection rates. KSHV is capable of infecting a broad range of cell types, including B cells, endothelial cells, epithelial cells, and fibroblasts (1). Like other herpesviruses, the KSHV life cycle comprises a lytic and latent viral program(2). Lytic infection results in the production of virus particles for intra- and inter-individual dissemination, while latency associates with the expression of key viral proteins (e.g. vCyclin; latency-associated nuclear antigen; LANA) that promote episomal persistence, host cell survival, and immune evasion, contributing to oncogenesis.

KS arises in four epidemiological forms: epidemic (associated with advanced HIV infection); iatrogenic (related to post-transplantation immunosuppression); endemic (occurring in Central and Eastern Africa); or classic (typically observed in the Mediterranean basin)(3). While the first two highlight the importance of host immunity in mitigating KSHV infection, KS onset, and/or severity, the latter two occur in immunocompetent hosts, where permissive immune defects remain undefined. Some rare KSHV infected individuals without know immunosuppression can develop endemic, or classic KS, mechanism remains unknown. However, therapy using topical or oral retinoid medication, or recombinant alpha interferon, have been shown to induce KS remission suggesting the involvement of two signaling pathways (4). Opportunistic infection without exogenous immunosuppression implies the presence of an intrinsic immunodeficiency. Supporting this premise, monogenic variants in *IFNGR1*, *WAS*, *STIM1*, *TNFRSF4 (OX40)*, *STAT4*, and *MAGT1*, genes predominantly implicated in T cell immunity, have been identified in rare patients with classic KS (cKS; table S1)(5–8). While these findings provide proof-of-concept that inborn errors of immunity may underlie susceptibility to cKS development, the mechanistic link between these variants and KSHV pathogenesis remains to be elucidated. Herein, we report a patient with classic KS due to autosomal recessive mutations in *DDX58* causing complete retinoic acid-inducible gene I (RIG-I) deficiency, uncovering a novel role for this cytosolic RNA sensor in human immunity to DNA viruses. RIG-I, a member of the RIG-I-like receptor (RLR) family (along with melanoma-differentiation-associated gene 5 (MDA5), and laboratory of genetics and physiology 2 (LGP2)(9)), typically detects short double-stranded RNA (dsRNA) molecules that have 5′-triphosphate or diphosphate ends, structures absent from host cytosolic RNA but commonly present in viral genomes or replication complexes. Upon ligand binding, RIG-I undergoes conformational changes that lead to its oligomerization through the recruitment of the mitochondrial adaptor MAVS, triggering downstream activation of TBK1 and IRF3/7, and culminating in the production of type I interferons(10). This pathway is critical in immunity to many RNA viruses, such as influenza A and hepatitis C virus(11–13).

Our work unveils RIG-I’s role in human immunity to KSHV, a DNA virus. We show, for the first time, that RIG-I critically mediates human anti-KSHV immunity, controlling both primary infection and reactivation from latency. Through transcriptomic and proteomic analyses, we demonstrate that RIG-I deficiency causes defective antiviral and inflammatory responses, promoting latent viral gene expression and oncogenic reprogramming. Given the mixed clinical results of IFN-α in the treatment of immunosuppression-associated KS (14, 15), we leveraged our findings to evaluate whether distinct type I interferon (IFN) subtypes could modulate KSHV latency in the context of impaired innate immune sensing (16, 17).

## RESULTS

### *DDX58* (RIG-I) mutation associated with Kaposi sarcoma

We investigated a man of Inuit origin (P1), from Salluit (Nunavik region in northernmost Quebec, Canada). He was initially referred at the age of 72 years for numerous violaceous nodules of the feet and pre-tibial regions. His past medical history included diabetes mellitus type 2, hypertension, dyslipidemia, coronary artery disease, and a transient ischemic attach in 2004. On initial assessment, and over the next 11 years prior to his death, he was repeatedly seronegative for HIV with no detectable HIV viral load. Histopathology revealed KS of the nodular type (Fig.1A). He was initially treated with liposomal doxorubicin for 2 cycles but stopped because of intolerance. Because of variable progression, resulting in chronic lymphedema of his right leg, he was administered cycles of paclitaxel at age 73, 75, 76, 77, 79, 81, and 83. At age 79, limited investigations for an underlying cause of his KS were pursued (tableS2) hampered by the geographic remoteness of his place of residence as well as the COVID-19 pandemic. He died at the age of 83 years from a cerebrovascular accident(18).

**Figure 1:**
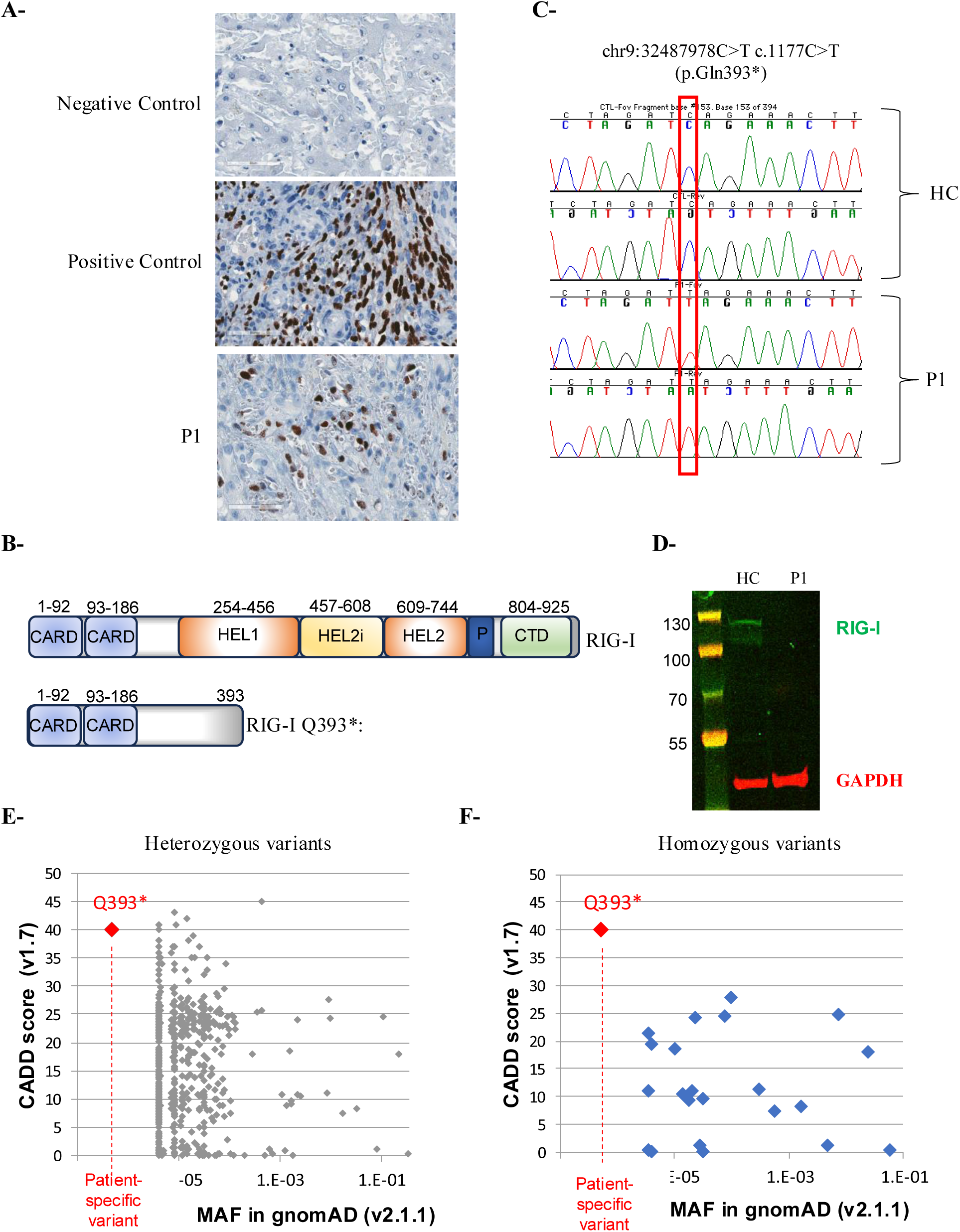
Clinical and molecular characterization of a patient with Kaposi Sarcoma. **A-** Immunohistochemistry for HHV-8 on a biopsy of violaceous nodule from P1 (negative and positive controls are indicated). **B-** Schematic representation of the RIG-I protein, highlighting the domain structure, including the CARD (Caspase Activation and Recruitment Domain), HEL (Helicase) domains, and CTD (C-terminal domain). The location of the p.Gln393* premature stop codon mutation is indicated. **C-** Sanger sequencing chromatograms confirming the homozygous c.1177C>T (p.Gln393*) mutation in the *DDX58* gene in P1. **D-** Western blot analysis of RIG-I protein expression in lymphoblastoid cell lines (LCLs) derived from a healthy control (HC) and the patient (P1). GAPDH serves as a loading control. The blot confirms the absence of RIG-I protein in P1 LCLs. **E-** F-Predicted CADD scores (https://cadd.gs.washington.edu/) and global allele frequencies of DDX58 variants (based on the gnomAD database (v2.1.1)). The red diamond indicates the variant of P1, whose CADD score was 40. The gray diamonds indicate heterozygous variants, and blue diamonds homozygous variants, respectively.

We hypothesized that the patient may have an underlying inborn error of immunity (IEI) as the cause for his susceptibility to KS. The extreme geographic isolation of the patient and presumably of his ancestors potentially suggested endogamy and thus, an autosomal recessive basis for the IEI. We performed whole-exome sequencing (WES), as previously done. We analyzed the data for variants in known IEI genes, including those previously reported with KS, and found none. Under a recessive model for rare (minor allele frequency ≤1:1000) and deleterious (e.g. CADD Phred ≥10) variants, 42 genes were identified (table S3). Of these, two were genes with immunity-related functions: c.1177C>T (p.Gln393*) in *DDX58* (encoding RIG-I) and c.314A>G (p.Asp105Gly) in *HLA-DRB1*. Given that *HLA-DRB1* is highly polymorphic in public (e.g. gnomAD) and private (e.g. Blacklist-Annotated variants) databases, we turned our attention to *DDX58*.

In humans, heterozygous missense mutations in *DDX58* resulting in molecular gain-of-function of RIG-I causes Singleton-Merten syndrome 2, characterized by glaucoma, aortic calcification, and skeletal abnormalities. In contrast, the chr9:32487978C>T variant in P1 was not found in public databases; it is a predicted premature stop codon (p.Gln393*), that in homozygosity, would lead to complete loss of expression (Fig. 1B). Notably, no disease with loss-of-function in *DDX58* has been reported. From genomic DNA of P1, we confirmed the homozygous variant by Sanger sequencing (Fig. 1C). From P1-derived lymphoblastoid cell lines (LCL), relative to healthy controls, no *DDX58* cDNA could be amplified by RT-PCR (data not shown), and no RIG-I protein was detected by Western blot using polyclonal antibodies (Fig. 1D), confirming that the identified c.1177C>T variant is null. We were not able to study the pedigree segregation of the *DDX58* variant, since there was no sample available from P1’s family members. *To better understand the potential impact of the p.Gln393* variant, we analyzed its distribution in relation to other *DDX58* variants catalogued in gnomAD. This variant was not documented in any public variant databases, including gnomAD v2.1.1, and no individuals carrying it in the homozygous state were identified. Furthermore, it displayed a CADD score of 40, indicating a high likelihood of deleteriousness and positioning it among the most damaging variants within the gene (Fig 1E and 1F). These findings implicate the RIG-I c.1177C>T (p.Gln393*) variant as a strong genetic candidate underlying P1’s KSHV susceptibility.

### *DDX58* p.Gln393* abrogates type I IFN responses to viral RNA agonists

To determine the functional consequences of RIG-I^Q393*^, we stimulated LCLs from P1 and healthy controls (HC) by transfection with the RIG-I-specific agonists 3p-hpRNA, poly(dA:dT), or 5’ppp-dsRNA and evaluated the expression of a panel of antiviral and antiproliferative interferon-stimulated genes (ISGs) (Fig. 2A). Response to non-RIG-I agonists (polyIC, R848, CpG ODN, 5ppp-CTL) were evaluated as controls. In P1’s LCL, ISG induction to all RIG-I-agonists was significantly blunted (Fig. 2A). Because RIG-I activates IRF3 to mount IFN responses, we quantitatively assessed RIG-I-dependent IRF3/type I IFN pathway activation using a ISG54-driven luciferase reporter A549-based cell line transfected with wild-type (RIG-I^WT^) or RIG-I^Q393*^, confirmed by Western blot (Fig. 2B). IFN-α treatment (serving as a RIG-I-independent positive control) triggered strong luciferase reporter activity in all conditions. Upon stimulation with 5’ppp-dsRNA (a RIG-I-specific agonist), RIG-I^WT^-expressing cells exhibited robust IRF3-pathway activation. On the other hand, RIG-I^Q393*^-expressing cells showed negligible IRF3-pathway activation, with responses comparable to empty vector-transfected controls (Fig. 2C). These results demonstrate that the RIG-I^Q393*^ mutation ablates IRF3-mediated type I IFN responses to viral RNA agonists, consistent with RIG-I’S canonical role in innate IRF3-mediated antiviral signaling.

**Figure 2:**
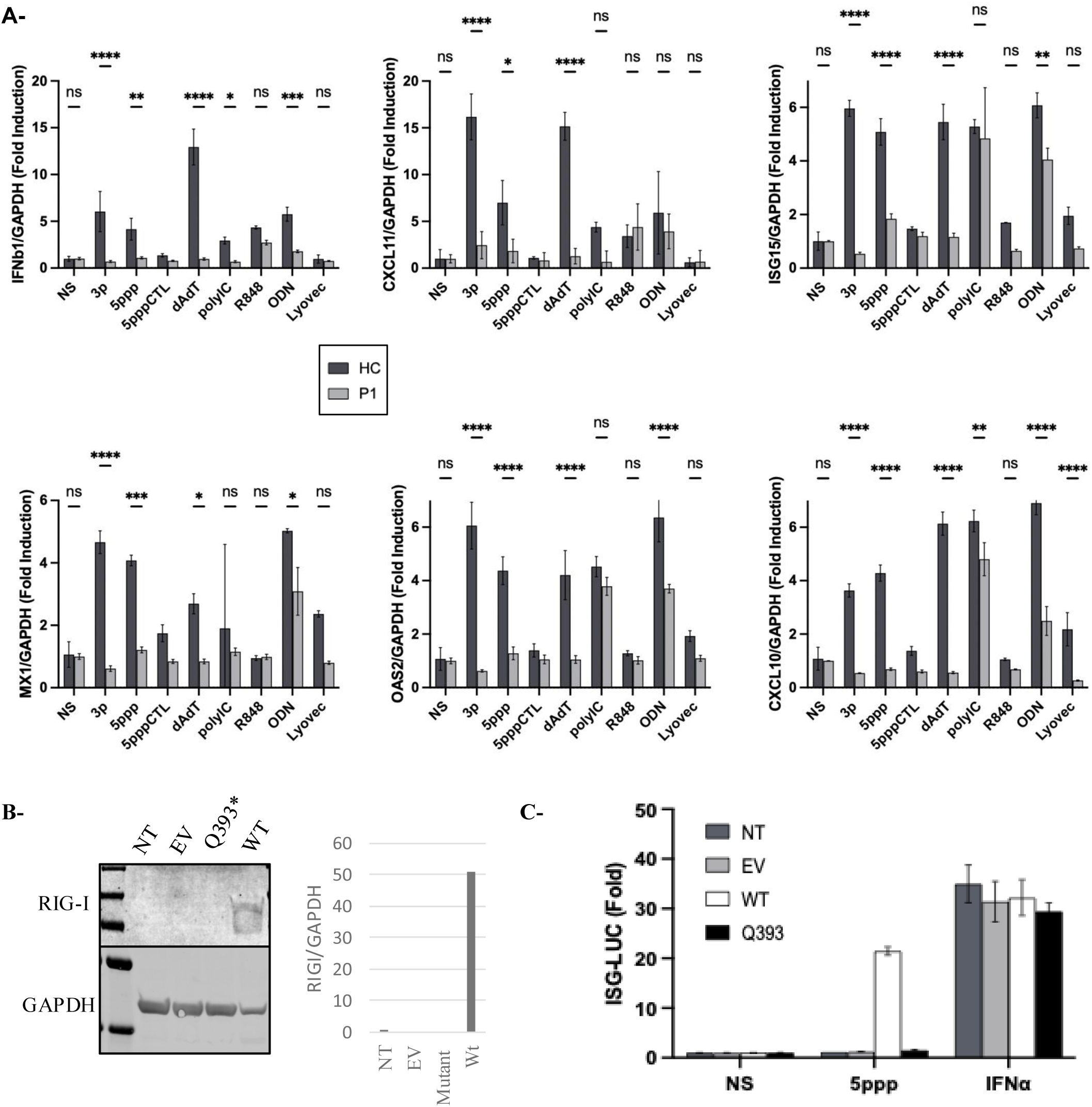
RIG-I deficiency impairs Type I Interferon responses to viral RNA agonists. **A-** LCLs from P1 and HC were stimulated for 24h with different RIG-I agonists and evaluated a panel of antiviral and antiproliferative interferon-stimulated gene (ISG) expression by QPCR. GAPDH is the housekeeping gene. **B, C-** A549-Dual™ KO-RIG-I cells were transfected with either an empty vector (EV), a wild-type RIG-I construct (RIG-I wt) or a RIG-I mutant construct (RIG-I Q393*). **B-** Immunoblotting was performed to evaluate RIG-I protein expression, and band intensities were quantified relative to GAPDH. **C-** Cells were then stimulated with 5’ppp or treated with IFNα (positive control) for 24h. Interferon regulatory factor (IRF) pathway activation was assessed by monitoring Lucia luciferase activity. Data are presented as relative luciferase units (RLU). Error bars represent standard deviation of the mean (SD) from triplicate experiments.

### Loss of RIG-I favors KSHV replication and skews the viral life cycle towards latency

While responses to molecular viral-RNA ligands are impaired by RIG-I^Q393*^, we sought to prove RIG-I’s relevance to host anti-KSHV immunity. We used iSLK-219 cells, a standard KSHV tool harboring KSVH.219 strain that bears dual fluorescent reporters (i.e. constitutive GFP to track infected cells, and doxycycline-inducible RFP to track lytic infection/reactivation) (fig. S1). We first used these iSLK-219-derived KSHV virions to infect the above A549/ISG54-cell line, showing that RIG-I deficiency also impairs IRF3-pathway activation during whole-virus infection (fig. S2). We also assessed RIG-I’s effect on KSHV dynamics by live-cell imaging (IncuCyte®), simultaneously tracking viral replication (KSHV-GFP) and host cell proliferation (Nuclight Red) for 72h in HUVECs with RIG-I either silenced by siRNA (siRIG-I) or treated with non-targeting siRNA controls (siCTL). siRIG-I cells exhibited increased GFP fluorescence relative to siCTL cells, indicating enhanced KSHV replication in the absence of RIG-I. Notably, siRIG-I HUVECs showed sustained cell proliferation following infection, in contrast to the reduced viability observed in siCTL cells. These results indicate that RIG-I is directly involved in immunity to KSHV: loss of RIG-I not only facilitates viral propagation but also disrupts infection-induced cell death. (Fig. 3).

**Figure 3:**
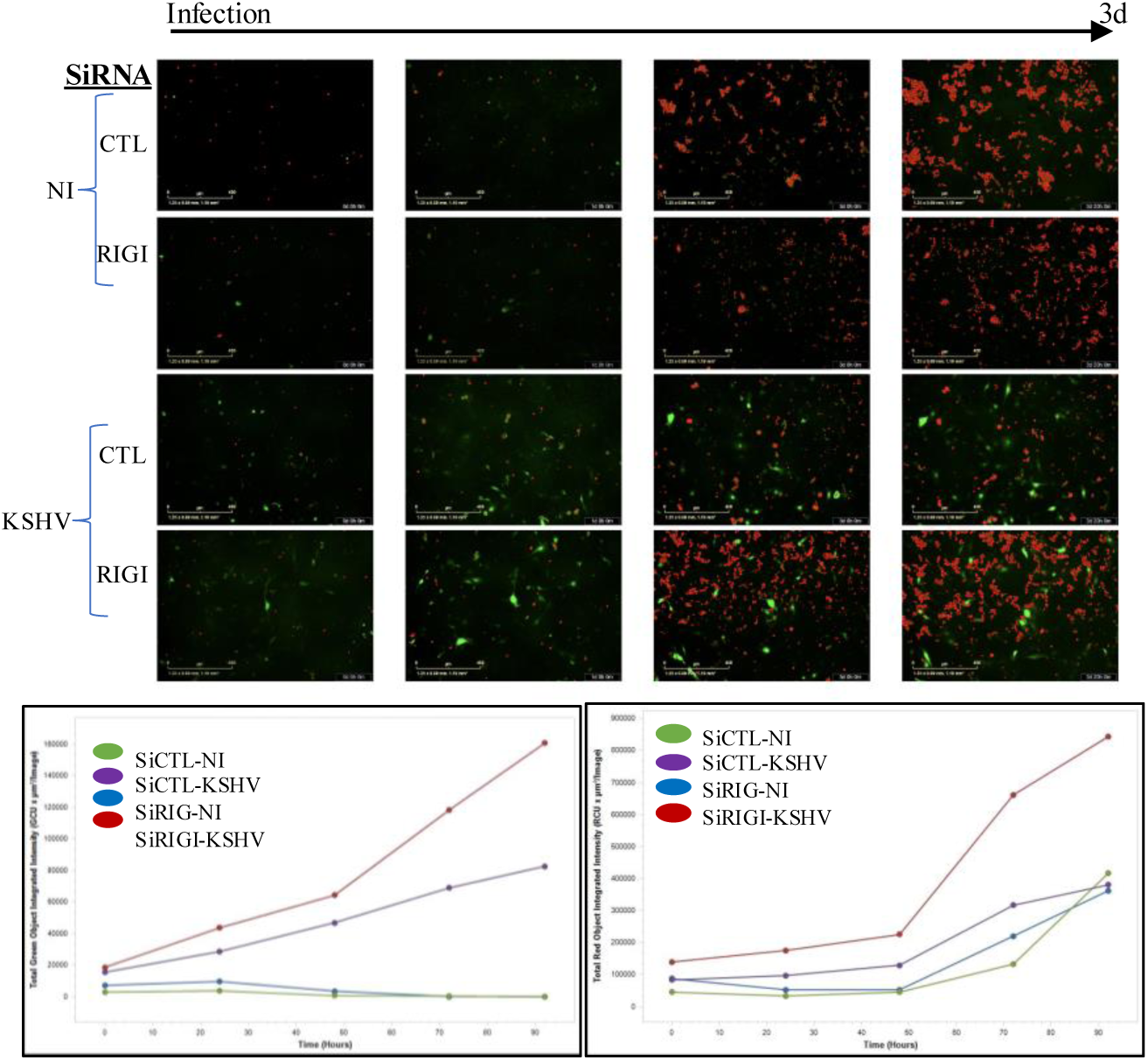
Representative IncuCyte images of HUVECs transfected with either siRNA CTL or siRNA siRIG-I, followed by infection with GFP-expressing KSHV. Images were acquired after 72 h using a 10× objective, showing NucLight Red stained nuclei (red) and GFP-positive KSHV-infected cells (green).

### Proteomic profiling reveals impaired inflammatory responses in RIG-I–deficient fibroblasts

To define how failed KSHV control during initial infection alters host-response outcomes beyond the acute infection phase, we performed targeted Olink proteomic analysis of supernatants collected three days post-infection of fibroblasts (RIG-I^WT^ and RIG-I^KO^). The RIG-I^KO^ fibroblasts were generated by CRISPR-Cas9 and RIG-I protein loss was confirmed by Western Blot prior to transcriptomic analysis (fig. S3). Global heatmap analysis (Fig. 4A) revealed broad alterations in normalized protein expression (NPX) values across all detected proteins in RIG-I^WT^ and RIG-I^KO^ fibroblasts. We observed significantly altered downregulation of 18 proteins (adjusted p>0.05, |log2FC|>1) in RIG-I^KO^ cells (Fig. 4B–D). Functional annotation (DAVID) revealed downregulation of antiproliferative cytokines LIF, IL-1α, and IL-24 (Gene Ontology term “negative regulation of cell population proliferation” (GO:0008285)), as well as of apoptotic regulators, including CASP8 and CXCL10 (GO:0006915) Genotype-specific clustering of these proteins was confirmed in a focused heatmap (Fig. 4D), and representative NPX distributions were visualized in boxplots (Fig. 4E), together demonstrating broad dysregulation of host defense pathways during intermediate-phase of infection.

**Figure 4.**
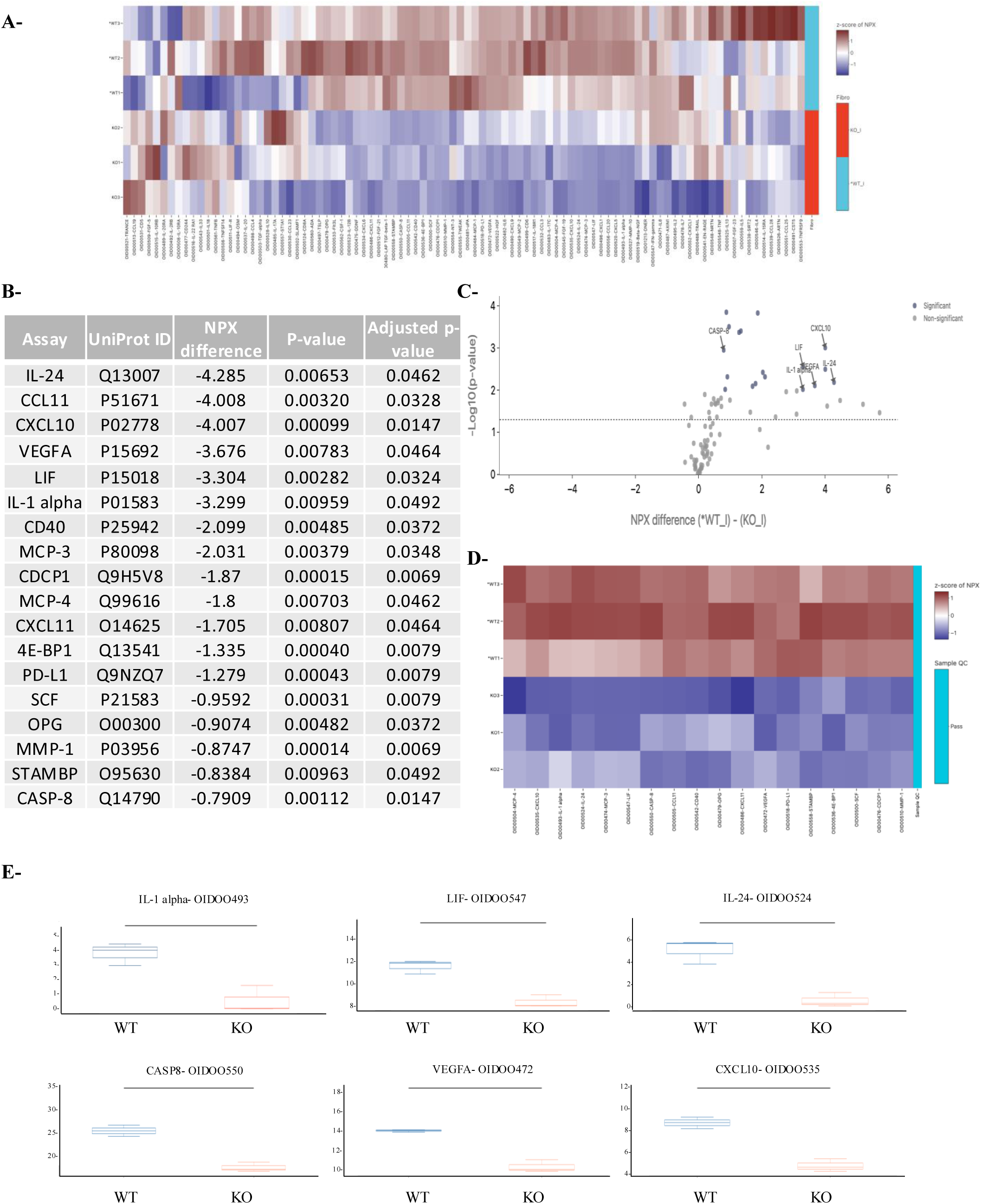
RIG-I deficiency alters the inflammatory secretome in KSHV-infected fibroblasts. WT and RIG-I KO fibroblasts (CRISPR-Cas9) were infected with KSHV for 72 h and cell culture supernatants were analyzed using the Olink® Target 96 Inflammation panel. **A-** Heatmap showing normalized protein expression (NPX) values across all detected proteins in WT and RIG-I KO fibroblasts. Samples are hierarchically clustered by genotype (blue = WT; red = KO). **B-** Table of the top 18 differentially expressed proteins between WT and RIG-I KO cells (Welch’s t-test, adjusted *p* < 0.05; Benjamini-Hochberg correction). **C-** Volcano plot showing NPX differences (x-axis) vs –log10(p-value) (y-axis). Labeled points highlight proteins passing the significance threshold (adjusted *p* < 0.05). **D-** Focused heatmap of the 18 significantly differentially expressed proteins from panel B. **E-** Boxplots displaying NPX distributions of selected top hits (*IL-24*, *LIF*, *IL-1*α, *VEGFA*, *CXCL10*, *CD40*) in WT versus RIG-I KO groups. Horizontal bars indicate median values. NPX = Normalized Protein eXpression (log_2_ scale). Statistical analysis was performed using the Olink® Statistical Analysis App v1.1.

### RIG-I loss remodels the transcriptional landscape in KSHV-infected fibroblasts

KSHV latency drives oncogenesis. To define the molecular mechanisms underlying this process, we established chronic KSHV infection in fibroblasts (RIG-I^WT^ and RIG-I^KO^) and assessed differential gene expression after one week. RNA sequencing (RNA-seq) analysis revealed genome-wide transcriptional perturbations in RIG-I^KO^ cells compared to RIG-I^WT^, with 483 differentially-expressed genes (FDR<0.05, |log2FC|>1) enriched in pathways associated with fibroblast activation, inflammation, and tumorigenesis (Fig. 5A: Heatmap, Fig. 5B: volcano plot). Gene ontology enrichment analysis of downregulated genes in RIG-I^KO^ fibroblasts revealed multiple significantly enriched biological processes, which were further organized into functionally related clusters using Metascape (Fig. 5C–D). Among the most strongly upregulated genes in RIG-I deficient fibroblasts were *VCAM1, POSTN, CCN4, HGF, and CXCL12*, which are functionally involved in extracellular matrix remodeling, angiogenesis, and proliferative signaling. These findings were validated through orthogonal approaches on independent biological replicates of KSHV-infected RIG-I^WT^ and RIG-I^KO^ fibroblasts: qPCR confirmed the significant upregulation of *VCAM1, POSTN, CCN4, HGF*, and *CXCL12* in RIG-I^KO^ cells (Fig. 5E), while Western blot demonstrated concordant increased VCAM-1 protein expression (Fig. 5F). These findings demonstrate that RIG-I deficiency results in transcriptional programs conducive to tumor microenvironment remodeling.

**Figure 5:**
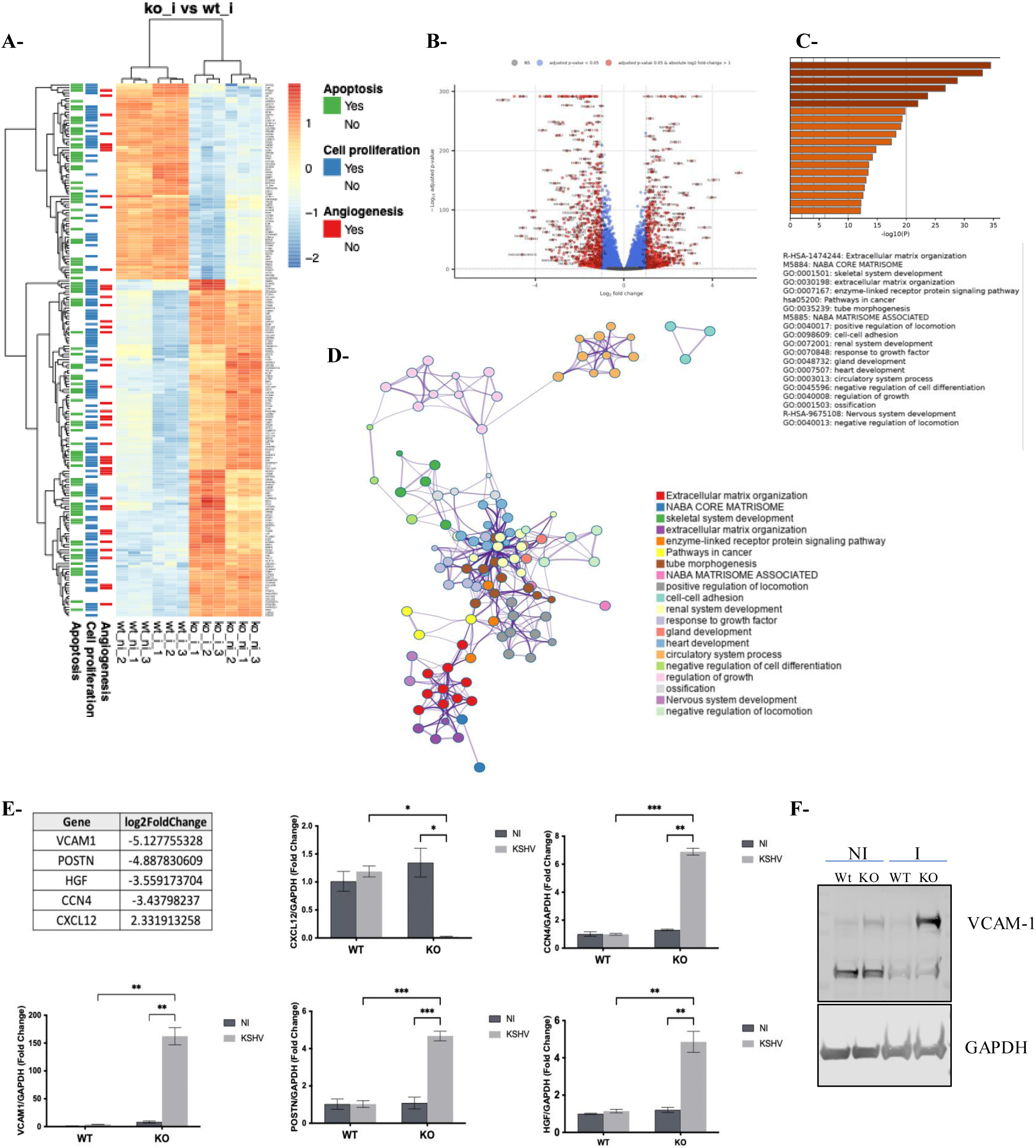
Loss of RIG-I compromises fibroblast control of KSHV. **A-** Heatmap of differentially expressed genes between ko_ni and ko_i related to apoptosis, cell proliferation and angiogenesis. The heatmap shows scaled expression values (z-scores) for selected DEGs across all samples. Genes were grouped by hierarchical clustering (rows), as were the samples (columns). The color scale represents relative expression levels: red indicates upregulation, and blue indicates downregulation. Functional annotations are shown as colored side bars for each gene, indicating known involvement in apoptosis (green), cell proliferation (blue), and angiogenesis (red). Each color indicates presence (“Yes”) or absence (“No”) of the gene in the corresponding enriched gene ontology term. **B-** Volcano plot of DEGs between KO_i and WT_i fibroblasts, highlighting genes with adjusted *p* < 0.05 and absolute log_ fold change > 1. **C-** Gene ontology enrichment analysis of downregulated genes in KO_i versus WT_i fibroblasts, showing the top 20 GO biological processes. **D-** Network visualization of enriched biological processes among DEGs, highlighting functionally related gene clusters. **E-** Validation by qPCR of five DEGs involved in angiogenesis and tumor microenvironment remodeling (VCAM1, POSTN, HGF, CCN4, and CXCL12) in WT and KO fibroblasts following KSHV infection. Relative expression levels are normalized to GAPDH and shown as fold change versus NI controls. Data represent means ± SD of triplicates. Statistical significance was determined using two-way ANOVA with Tukey’s post hoc test (*p* < 0.05, \**p* < 0.01, \*\**p* < 0.001). **F-** Western blot analysis of VCAM-1 expression in WT and RIG-I KO fibroblasts, either non-infected (NI) or infected (KSHV) for 7 days. GAPDH was used as a loading control.

### Absence of RIG-I skews primary infection towards KSHV latency

To delineate the viral mechanisms driving these dysregulated host cell responses, we infected fibroblasts with KSHV and observed distinct latency profiles. Compared to RIG-I^WT^ controls, RIG-I^KO^ fibroblasts showed significantly increased expression of key latent viral genes *LANA* and *vCyclin* (Fig. 6A), with only modest reduction in the lytic gene vIL-6. Moreover, in RIG-I^KO^ cells, we observed marked upregulation of the human circular RNA, hsa_circ_0001400, that was previously reported to promote latency and cell cycling, suppressing lytic replication and apoptosis(19–21), suggesting RIG-I maintains viral latency equilibrium through both direct and RNA-mediated pathways. Western blot analysis confirmed increased levels of LANA protein in RIG-I^KO^ cells (Fig. 6B). To validate these results in a cell-independent context, we repeated the experiments in human microvascular endothelial cells (HMEC-1), another physiologically-relevant target of KSHV. Similar to fibroblasts, RIG-I knockdown in HMEC-1 resulted in elevated expression of latent genes (*vCyclin*) and *hsa_circ_0001400* (fig. S4), supporting a conserved role for RIG-I in restraining KSHV latency across different cell types.

**Figure 6:**
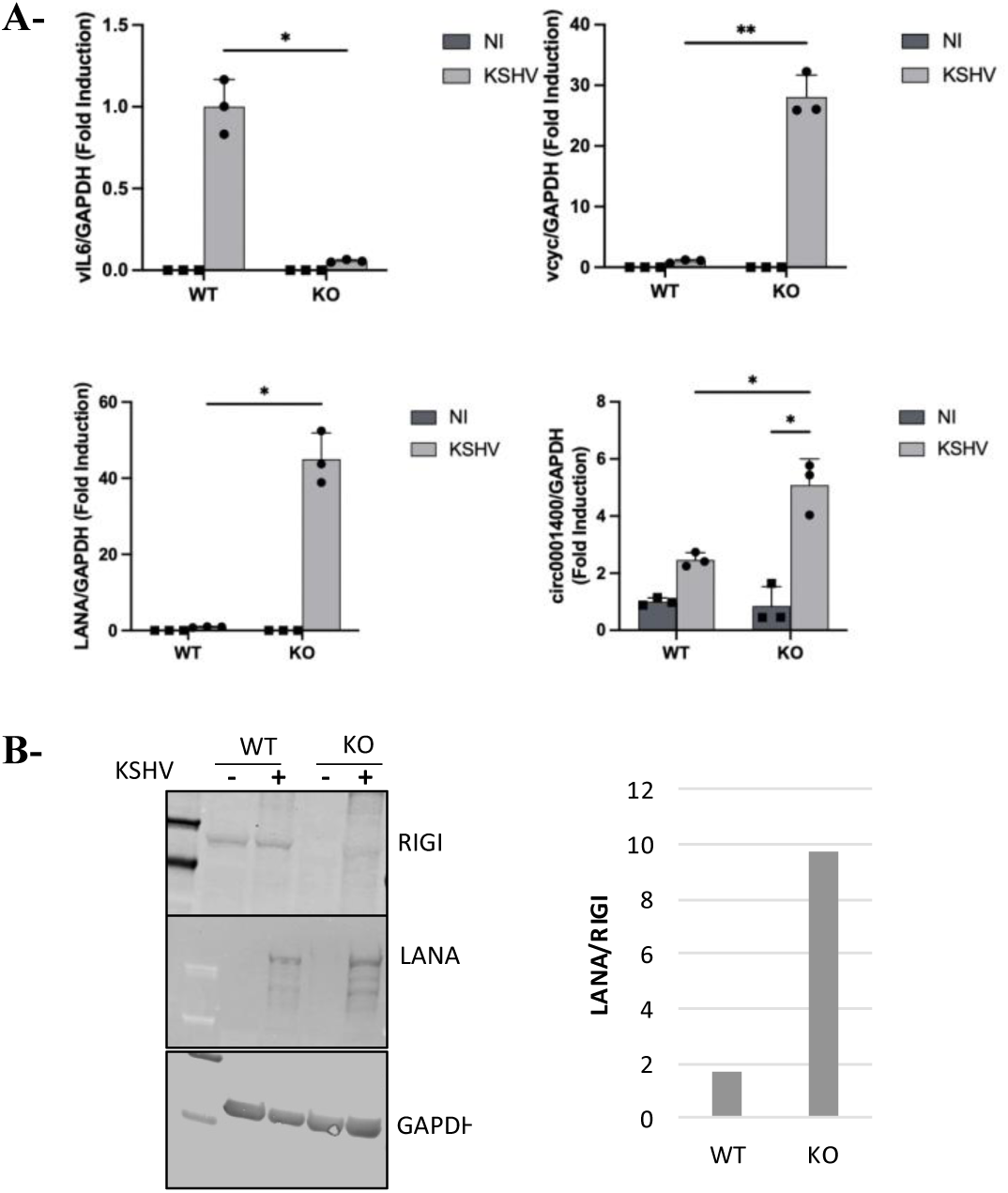
Loss of RIG-I promotes KSHV replication and skews the viral life cycle toward latency. IMR-90 RIG-I knockout fibroblasts were infected with KSHV for 7 days. **A-** Total RNA was extracted and analyzed by RT–qPCR for KSHV latent (*LANA*, *vCyclin*) and lytic (*vIL-6*) gene expression, as well as the host circular RNA *hsa_circ_0001400*. Data are presented as fold change relative to uninfected wild-type (WT NI) controls and normalized to *GAPDH*. Error bars indicate the standard deviation from triplicate experiments. **B-** Western blot analysis of LANA and RIG-I protein levels in WT and RIG-I knockout fibroblasts infected with KSHV for 7 days. GAPDH was used as a loading control.

### RIG-I constrains KSHV reactivation, host cell remodeling, and oncogenic program

Given our findings that RIG-I is critical in host defense to primary infection, we examined its function in regulating responses during the latent-to-lytic switch in KSHV infection (reactivation). Using the iSLK.219 cell line, which is fully permissive for doxycycline-induced, complete KSHV reactivation to the lytic program, we examined the impact of RIG-I knockdown. Silencing of RIG-I led to a dramatic increase in expression of the lytic gene *vIL-6, accompanied by a moderate but also significant elevation of* the latent genes, *LANA* and *vCyclin*, indicating RIG-I’s role in constraining viral reactivation (Fig. 7A). To further define the impact of RIG-I on KSHV reactivation at the cellular level, we analyzed primary HUVECs following RIG-I knockdown, KSHV infection, and treatment with the histone deacetylase inhibitors, trichostatin A (TSA) and sodium butyrate (NaB), which induce chromatin remodeling to suppress latent gene expression and chemically induce reactivation(22, 23) (Fig. 7B). Immunofluorescence analysis revealed RIG-I’s role in regulating cellular actin responses: RIG-I-expressing cells exhibited a quiescent cytoskeletal phenotype. In contrast, siRIG-I cells developed extensive actin network reorganization, indicating enhanced cytoskeletal remodeling and cellular activation. These results position RIG-I as a nexus linking innate immunity to both KSHV latency control and tumor-associated cytoskeletal dysregulation. Further corroborating RIG-I’s regulatory role, we observed that the elevated baseline LANA levels seen in KSHV-infected fibroblasts with RIG-I depletion were reduced upon TSA/NaB-induced reactivation, consistent with a shift towards a lytic viral program (Fig. 7C). This reduction in LANA expression correlated with diminished expression of top differentially-expressed host genes identified by RNA-seq, supporting a model wherein LANA orchestrates transcriptional programs that drive cellular transformation. Altogether, these findings establish RIG-I as a dual-function regulatory in KSHV infection: it not only controls primary infection, but it also critically regulates host responses to viral reactivation, modulating cellular remodeling and oncogenic transcriptional programs.

**Figure 7:**
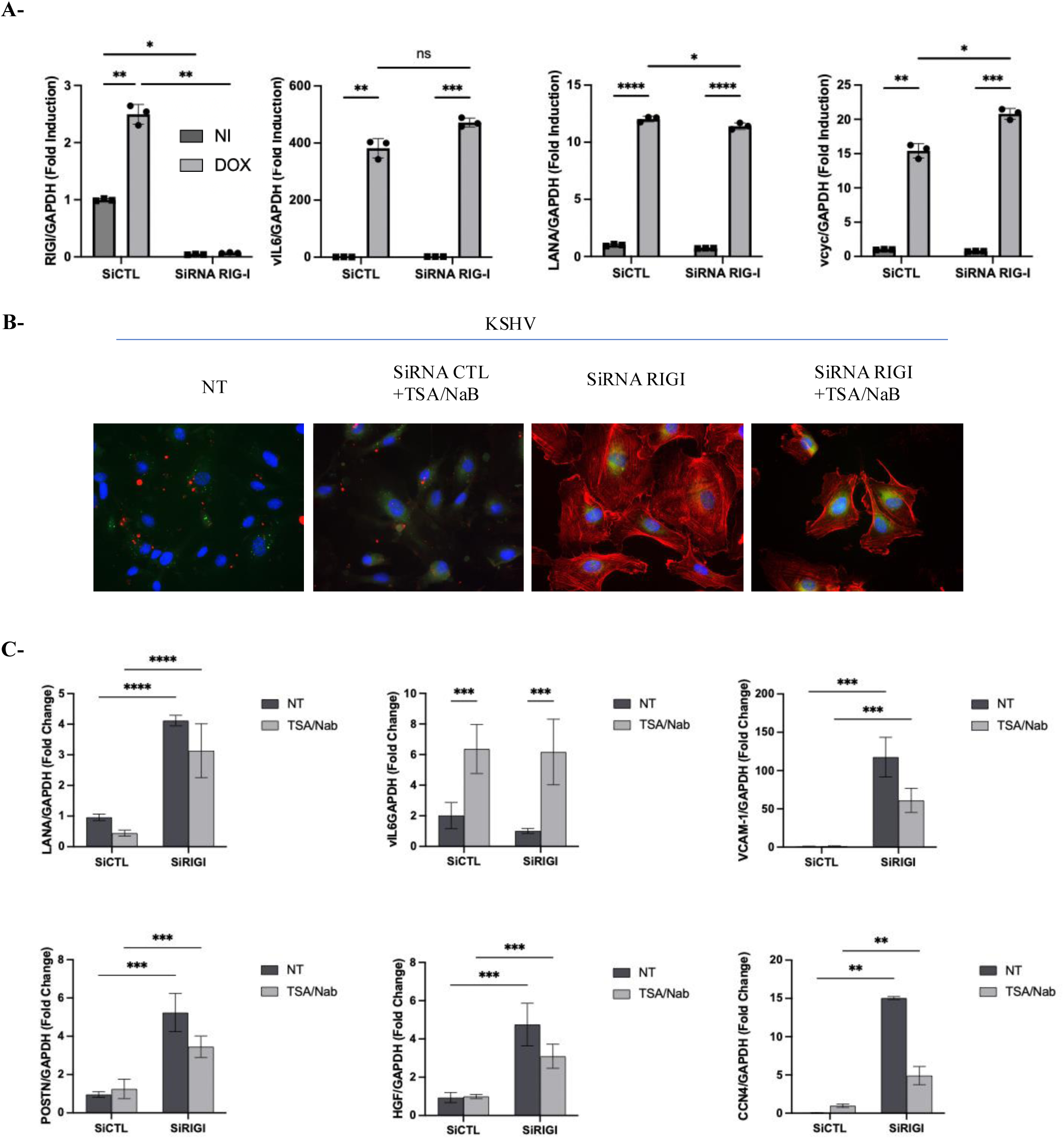
RIG-I deficiency alters viral gene expression and cytoskeletal structure following KSHV infection. **A-** iSLK.219 cells harboring latent KSHV were transfected with siRNA CTL or siRNA RIG-I. At 24 h post-transfection, lytic reactivation was induced with doxycycline (Dox) for 48 h. Total RNA was extracted and analyzed by RT–qPCR for KSHV transcript levels. **B-** HUVECs were transfected with siRNA CTL or siRNA RIG-I and infected with KSHV under the same conditions as in (A). Cells were subsequently treated with TSA/NaB for 24 h, fixed, and stained to visualize actin filaments (phalloidin, red), nuclei (DAPI, blue), and KSHV markers (green). Images were acquired at ×40 magnification. **C-** IMR-90 fibroblasts were transfected with control siRNA CTL or siRNA RIG-I. After 24 h, cells were infected with KSHV (MOI 1). At 24 h post-infection, cells were treated with sodium butyrate (NaB, 1 mM) and trichostatin A (TSA, 500 ng/mL) for 24 h. Total RNA was extracted with TRIzol and subjected to RT–qPCR to assess viral gene expression.

### Type I interferons partially restore antiviral control of KSHV in RIG-I–deficient cells

To assess whether exogenous IFN administration could restore antiviral control in the absence of RIG-I, we infected fibroblasts (Fig. 8A) and A549 cells (Fig. 8B) with KSHV, comparing RIG-I^WT^ to RIG-I^KO^ for each cell type. Seven days post-infection, cells were treated for 24 hours with different type I IFN subtypes (IFN-α, IFN-β, IFN-ω; each at 1000 IU/mL), and the expression of the latent viral genes *LANA* and *vCyclin* was quantified by RT-qPCR. In both cell types, under unstimulated conditions (NS), RIG-I^KO^ cells exhibited significantly higher expression of *LANA* and *vCyclin* compared to their WT counterparts, confirming our above findings on the role of RIG-I in the basal control of viral latency. Treatment with IFN-α led to a modest reduction in *LANA* and *vCyclin* expression in RIG-I^KO^ cells after KSHV infection. Strikingly, IFN-β and IFN-ω treatment resulted in a marked decrease of these latent genes in RIG-I-deficient cells, reaching levels comparable to those observed in treated WT cells. To corroborate these findings, we used KSHV-GFP–infected A549 cells and monitored latent viral expression by flow cytometry (Fig. 8C). Consistent with the RT-PCR results, IFN-β and IFN-ω treatment significantly reduced GFP fluorescence intensity in both WT and RIG-I^KO cells, indicating suppression of the latent viral program. These results demonstrate that the functional paralysis that occurs with RIG-I deficiency towards KSHV can be rescued with type I IFN treatment.

**Figure 8:**
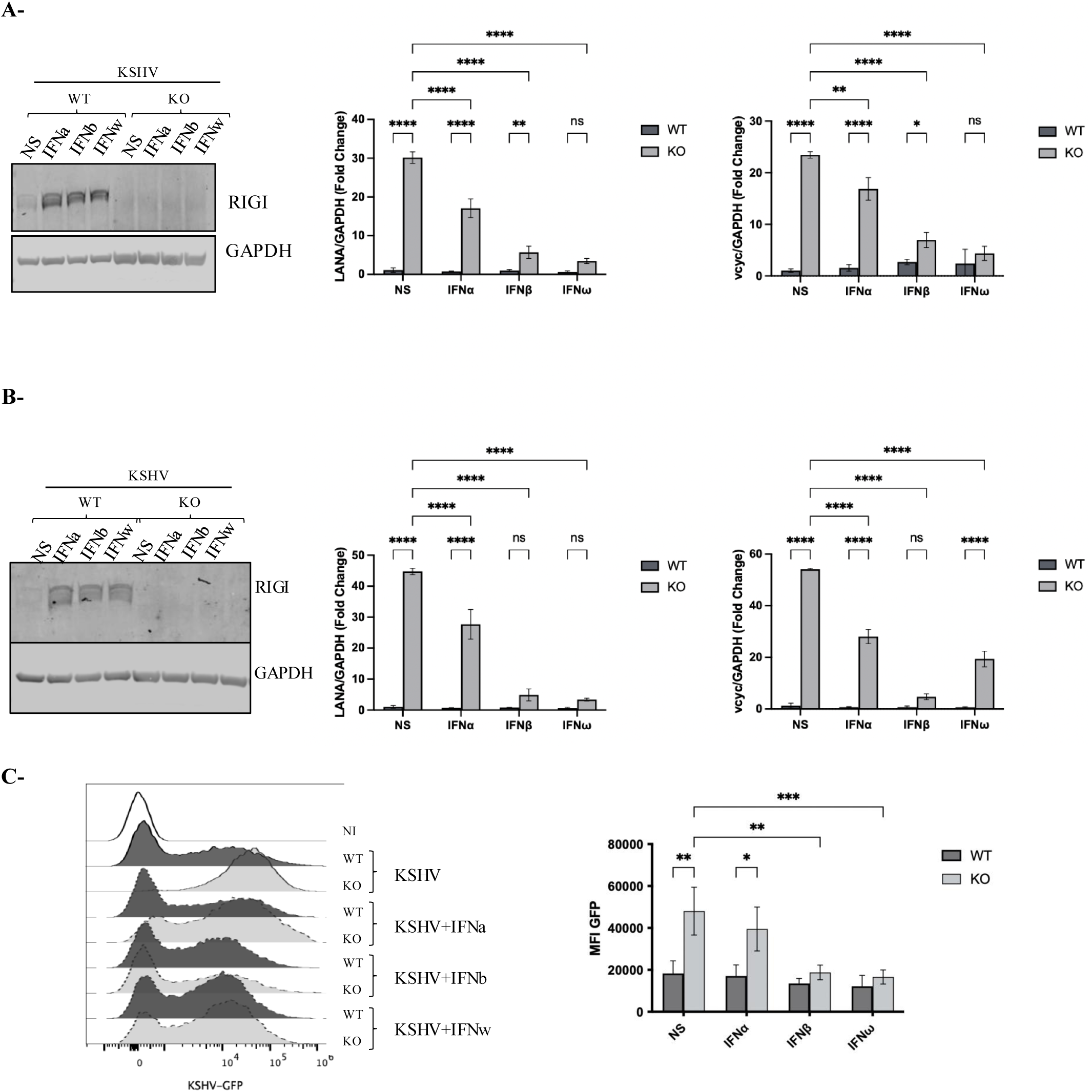
Type I IFN treatment suppresses KSHV latent gene expression in RIG-I–deficient cells. **A-** IMR-90 fibroblasts and **B-** A549 epithelial cells (RIG-I^wt^ and RIG-I^KO^) were infected with KSHV (MOI = 1). After 7 days, cells were treated for 24 hours with type I interferons (IFN-α, IFN-β, or IFN-ω; 1,000 IU/mL). Left panels: immunoblot confirming RIG-I deficiency in CRISPR-Cas9–edited fibroblasts and RIG-I^KO^A549 cells. GAPDH serves as a loading control. Right panels: quantitative RT–PCR analysis of latent KSHV transcripts (LANA and vCyclin) normalized to GAPDH. **C-** A549 cells (WT and RIG-I KO) were infected with KSHV-GFP and treated 7 days later with IFN-α, IFN-β, or IFN-ω (1,000 IU/mL) for 24 h. Left: representative flow cytometry histograms of GFP expression; dotted lines represent IFN treatments. Right: mean fluorescence intensity (MFI) of GFP expression (KSHV-GFP). Data are presented as mean ± s.d. of biological replicates. Statistical significance was determined by two-way ANOVA with multiple comparisons; *p<0.05, **p<0.01, ***p<0.001, ****p<0.0001; ns, not significant.

## DISCUSSION

Classic KS corresponds to the original description by Moritz Kaposi in 1872, in which reddish-purple nodules or plaques appear, typically in middle or older aged male adults, first on the feet and lower extremities, then the hands, and may progress indolently (over years) to widespread involvement. While it has been primarily associated with individuals of Mediterranean, Eastern European or Ashkenazi Jewish origin, there have been sporadic reports of classic KS in other ancestral backgrounds, including the Uyghur in Xinjiang, China(24); Indigenous groups in Peru(25) and Columbia(26); and the circumpolar Iñupiat populations, i.e. the Inuit of Canada, Greenland, and northern Alaska(27–32). The basis for these classic KS cases is not well understood, but given the historical endogamy of some peoples, a genetic basis is suspected. Herein, we have identified an inborn error of innate antiviral immunity underlying classic KS in an elderly male of Inuit origin. Whether this finding implies genetic lesions of these pathways in other affected individuals of the Nunavik arctic region, or in other Inuit persons of northern Canada or Greenland, requires appropriate partnership with the respective communities to evaluate further.

Our study reinforces and extends prior mechanistic understanding of RIG-I involvement during KSHV infection. Although previous studies have provided important insights into the molecular mechanisms of RIG-I activation, they largely relied on ectopic systems using synthetic or virus-derived RNA fragments. These approaches demonstrated that RIG-I can sense misprocessed host RNAs such as 5′-triphosphorylated RNAs accumulating upon DUSP11 downregulation (33, 34), or structured KSHV transcripts through a RNA polymerase III-independent mechanism (33, 34). Our study complements these data by showing that in physiologically relevant structural cells (fibroblasts and endothelial cells) supporting the full KSHV life cycle, RIG-I plays an active role in antiviral defense by promoting IRF3 activation and type I interferon responses. Thus, beyond sensing immunostimulatory RNA species, RIG-I emerges as a critical antiviral effector operating in the context of an oncogenic DNA virus, underscoring its broader role in host defense beyond RNA viruses.

To further delineate the consequences of RIG-I deficiency, we conducted transcriptomic analyses of RIG-I-deficient fibroblasts infected with KSHV. One-week post-infection, RNA-seq revealed the upregulation of genes involved in key tumor-promoting processes such as cell proliferation, migration, and microenvironment remodeling. Notably, genes such as VCAM-1, CCN4 (WISP) and HGF were consistently elevated. These genes converge on shared oncogenic functions. Specifically, VCAM-1 is often secreted by cancer-associated fibroblasts, thereby supporting tumor growth and invasiveness via activation of AKT and MAPK pathways (35). CCN4 encodes a matricellular protein regulated by Wnt signaling that facilitates cancer cell proliferation, migration, and extracellular matrix remodeling (36). HGF, a mesenchymal-derived growth factor, promotes epithelial-to-mesenchymal transition (EMT) and vasculogenic mimicry, both linked to enhanced invasiveness and metastatic potential in hepatocellular carcinoma and other cancers (37). Collectively, the transcriptional upregulation of these genes in RIG-I-deficient cells supports the notion that loss of antiviral surveillance in the context of persistent viral infection may drive a pro-tumoral phenotype, underscoring the role of RIG-I in restraining oncogenic programs activated by KSHV.

In parallel, we evaluated the early host response to KSHV infection through proteomic analyses of fibroblast supernatants (RIG-I^WT^ or RIG-I^KO^) using the Olink Inflammation panel. In RIG-I-deficient cells, several cytokines with anti-proliferative and tumor-suppressive functions including IL-24, CASP8, and CXCL10 (38–40) were statistically downregulated. These cytokines have been shown to inhibit cell proliferation and promote apoptosis of cancerous cells. The combined decrease of these main mediators in the absence of RIG-I reflects a shift toward a pro-tumoral immune environment, even during acute infection. Notably, while these proteomic data imply that RIG-I helps maintain a tumor-suppressive state early in KSHV infection, the lack of a broader pro-tumoral cytokine signature is consistent with established models where pro-oncogenic immune remodeling emerges predominantly during chronic, rather than acute, infection. Chronic infections are associated with sustained inflammation, immune evasion, and tissue remodeling that facilitate tumor progression, whereas acute infections tend to be transient and may even exert protective, anti-tumor effects through robust immune activation (41). Our results suggest that RIG-I deficiency licenses tumor-promoting transcriptional programs earlier in the infection cycle compared to RIG-I-sufficient cells, implicating a breakdown in protective mechanisms that normally constrain oncogenesis during the initial infection phases.

Importantly, we observed increased expression of KSHV latent genes, *LANA* and *vCyclin*, in RIG-I-deficient cells. These latency-associated proteins are known to drive oncogenic transformation: LANA promotes chromosomal instability by targeting the mitotic checkpoint kinase Bub1 for degradation (42–44), while vCyclin accelerates cell cycle progression and impairs pRB-mediated tumor suppression(45). Our findings are corroborated by recent studies showing that the circular RNA *circ_0001400*, induced during KSHV infection, enforces latency and supports immune evasion and cell survival(46). Taken together, these data define RIG-I as a gatekeeper not only for viral clearance but also for preventing viral-driven oncogenesis.

To further delineate the role of RIG-I during distinct phases of the KSHV life cycle, we employed complementary cellular models that recapitulate both primary infection and viral reactivation. Having established the importance of RIG-I during primary infection in fibroblasts, we next confirmed its involvement in the reactivation phase. The iSLK.219 cell line, which harbors latent KSHV and allows for synchronous induction of the lytic cycle, enabled us to examine viral reactivation dynamics. Our data demonstrate that RIG-I has dual sentinel roles restricting both primary infection and latent-to-lytic reactivation.

Results from our study also highlight the functional importance of RIG-I in restricting KSHV latency and reveal that exogenous cytokine therapy can rescue, at least partially, this antiviral function in RIG-I-deficient settings in vitro. In this context, we comprehensively assessed whether distinct type I interferon subtypes differentially modulate the viral latency program. We found that treatment of KSHV-infected cells with IFN-β or IFN-ω led to a marked reduction in *LANA* and *vCyclin* latent gene expression, with IFN-α exerting only a modest effect. These results align with prior studies demonstrating that IFN-β and IFN-ω elicit broader and more sustained antiviral responses compared to IFN-α, including enhanced ISG induction and chromatin remodeling (16, 17). The superior antiviral efficacy of these subtypes against other DNA viruses, such as HSV-1 and CMV, as reported by Lin et al., corroborates our observations in the context of KSHV (16). Similarly, the limited capacity of IFN-α to suppress viral latency, as shown by Zhang et al., mirrors the partial effect we observed on KSHV latent transcripts (17). Although IFN-α remains widely used in antiviral therapies, its suboptimal effectiveness against DNA viruses like KSHV, along with its unfavorable tolerability profile (14, 15), underscore the need to explore alternative strategies. Our results suggest that IFN-β and IFN-ω may represent more effective therapeutic options, particularly in the setting of innate immune deficiencies such as loss of the RIG-I-associated pathway. While the dose we used in vitro (1000 IU/mL) is a common research dose in cell cultures, it exceeds typical serum levels achieved clinically(47). Nonetheless these *in vitro* experiments support the concept of subtype-specific interferon therapies as a rational approach to target KSHV latency and warrant further clinical investigations.

In summary, we describe the first IEI in innate antiviral immunity conferring susceptibility to KSHV, causing classic KS. This work expands RIG-I’s role in human antiviral immunity beyond canonical RNA virus infections, advances KSHV immunopathogenesis, and offers therapeutic targets.

## MATERIALS AND METHODS

### Human subjects

Written informed consent was obtained by all patients or their legally authorized representative. The study was approved by the McGill University Health Centre Research Ethics Board protocol 10-256.

### Sequencing and bioinformatics analysis

For Sanger sequencing, the *RIG-I/DDX58* gene was PCR amplified from genomic DNA using primers designed to flank the respective regions (primers and sequencing conditions available on request). Sequencing was performed at the *Génome Québec* Innovation Centre (Montreal, Canada). Sequencing analyses were performed on Sequencher® sequence analysis software (Gene Codes Corporation, Ann Arbor, MI).

### Cell culture

Fibroblasts (IMR-90) were purchased from ATCC and were cultured in EMEM supplemented with 10% fetal bovine serum (FBS) and penicillin/streptomycin (100U/ml and 100ug/ml respectively; Wisent). Immortalized human dermal microvascular endothelium HMEC-1 cells were purchased from ATCC CRL-3243 (Rockville, MD, USA) and cultured in MCDB 131 medium containing 10ng/mL Epidermal Growth Factor (EGF), 1 µg/mL Hydrocortisone, 10 mM Glutamine, 10% fetal bovine serum. EBV-induced lymphoblastoid cell lines (LCL) were established, as previously described(Vinh et al., 2011, Gavino et al., 2014, Gavino et al., 2016). LCLs were cultured in RPMI supplemented with 10% FBS, 20mM HEPES and antibiotics. HUVECs were grown in 199 medium, supplemented with 20% heat-inactivated FBS, 60 μg/ml endothelial cell growth supplement, 2 mM L-glutamine, and 50 μg/ml heparin. HUVECs were cultured from passages 8. A549-Dual™ KO-RIG-I were purchased from InvivoGen (San Diego, CA) and were grown in Dulbecco’s modified Eagle’s medium (DMEM; Life Technologies) supplemented with 10% FBS, Normocin™ (100 μg/ml), blasticidin (10 μg/ml) and Zeocin™ (100 μg/ml). iSLK.219 cells (a kind gift from Dr. Arias) were maintained in DMEM medium (Corning) containing 10% tetracycline (Tet)-free FBS, 1% Pen-Strep, 10 μg/ml puromycin (Corning), 250 μg/ml G418 and 400 μg/ml hygromycin B(48, 49).

### Antibodies and stimulations

RIG-I polyclonal antibody was from Invitrogen. Anti-GAPDH was from Millipore. Secondary antibodies for immunoblotting: DyLight800-anti-rabbit-IgG and DyLight680-anti-mouse-IgG were from ThermoScientific. were from Cell Signaling Technology. 3p-hpRNA, 5’ppp-dsRNA, 5’ppp-dsRNA Control, Poly(dA:dT), Poly(I:C), R848, ODN 2006, Poly(dA:dT) and Poly(dG:dC) and Lyovec were from Invivogen. IFN-alpha was from PBL Assay Science. IFN-alpha and IFN-beta were from PBL Assay Science. IFN-omega was from ebioscience. VCAM-1 antibody was from Invitrogen. Incucyte Nuclight Red Dye was from Sartorius. Trichostatin A (TSA) and sodium butyrate (NaB) were from MilliporeSigma.

### Quantitative real-time PCR

RNA from cells was extracted using Trizol Reagent (Invitrogen) per manufacturer’s instructions and quantified by NanoDrop-100 spectrophotometer. 500ng of RNA was reverse transcribed using the Maxima H Minus First Strand cDNA Synthesis Kit (Thermo Fisher Scientific). Real-time quantitative PCR was performed with CFX Opus 96 Real-Time PCR (Bio-Rad Laboratories). TaqMan quantitative PCR assays were run using primers/probes: *IFNb* (Hs 01077958), *CXCl11* (Hs00171138), *CXCL10* (Hs00171042), *OAS2* (Hs00942643), *MX1* (Hs00895608), *CXCL12* (Hs 03676656), *CCN4* (Hs 05047584), *POSTN* (Hs 1566750), HGF (Hs 1566750) and RIG-I (Hs 01061436). The mRNA input was normalized to the expression of the housekeeping gene, *GAPDH* (Hs02786624). For SYBR Green-based detection, the sequences of the primer pairs are vIL-6 (F:5’-TTCAAAACACGCACCGCTTG-3’, R:5’-AAACGTGGACGTCATGGAGC-3’), LANA (F:5’-TTTAGTGTAGAGGGACCTTGGG-3’, R:5’-TCTCCATCTCCTGCATTGCC-3’), vcyc (F:5’-CGCCTGTAGAACGGAAACAT-3’, R:5’-TTGCCCGCCTCTATTATCAG-3’), circ0001400 (F:5’-ATGTCTGTTAGTGGGGCTGA-3’, R:5’-TATCTGCTACCATCGCCTTT-3’), GAPDH (F:5’-GTCATCAATGGAAATCCCATCACC-3’, R:5’-TGAGTCCTTCCACGATACCAAA-3’).

### Viruses and infection

KSHV inoculum was generated from iSLK.219 cells induced with 0.2 µg/mL doxycycline (DOX) in antibiotic-free medium. After 5 days, culture supernatants were harvested, clarified by centrifugation (1,000 × g for 10 min at 4_°C), and filtered through a 0.45-µm pore-size cellulose acetate filter (Corning). After centrifugation, concentrated virions were either used immediately or stored at –80_°C. Viral titration was performed by quantitative PCR using the Genesig Easy kit for Human Herpesvirus 8. Cells were infected at a multiplicity of infection (MOI) of 1 in the presence of 8_µg/mL polybrene (Sigma) by spinoculation (centrifugation at 1,500 × g for 60 min). After infection, the medium was replaced with fresh culture medium and cells were incubated at 37_°C for the indicated duration.

### DNA constructs and transfection

The plasmid encoding wild-type (WT) RIG-I was generated using the pCMV6-myc-DDK vector (OriGene, PS100001). Site-directed mutagenesis (New England Biolabs) was used to introduce the variant of interest into the WT construct. Cells were transfected with plasmid DNA using Lipofectamine 3000 (Invitrogen), following the manufacturer’s protocol. For gene silencing, RIG-I–targeting siRNA (SiRNA RIG-I) or non-targeting control siRNA (SiRNA CTL) (Invitrogen) were transfected using Lipofectamine RNAiMAX (Invitrogen) for either 48 or 72 hours, depending on the experiment.

### Immunoblotting

Cells were resuspended in RIPA lysis buffer (Sigma) supplemented with cOmplete Mini Protease Inhibitor Cocktail and PhosSTOP (both from Roche), then boiled in LDS Sample buffer and Blot Sample Reducing Agent (Thermo Fisher Scientific). Protein was separated by SDS-PAGE on premade gels (Novex) before being transferred to a membrane via iBlot Gel Transfer Device (Invitrogen). Membrane was blocked in 5% skim milk TBST to block nonspecific protein binding. The membrane was blotted using primary antibodies overnight at 4°C. The membrane was washed 4 times with 0.1% Tween 20 in TBS and then incubated with a 1:15,000 dilution of secondary antibodies in blocking buffer. Membranes were scanned and analyzed with an Odyssey IR scanner using Odyssey imaging software 3.0 (LI-COR Biosciences, Inc).

### IRF reporter assay

Activation of the interferon regulatory factors (IRF) pathway was monitored by measuring Lucia luciferase activity in A549-Dual™ RIG-I^−/−^ cells. It was evaluated using QUANTI-Luc substrate according to manufacturer’s recommendations. Briefly, 100 µL of QUANTI-Luc substrate was added to 20 µL of cell culture supernatant, and IRF-induced luciferase levels were quantified by luminescence measurement using a plate reader (Tecan M1000). Results are presented as fold-change ratio of luminescence emitted by supernatant from treated cells in comparison with control cells.

### Live-cell imaging of KSHV infection in siRNA-transfected HUVECs

Human umbilical vein endothelial cells (HUVECs) were seeded in 48-well plates and transfected with either control siRNA CTL or siRNA RIG-I. At 24 h post-transfection, cells were infected with GFP-expressing KSHV in the presence of 8 μg/mL polybrene by spinoculation. NucLight Rapid Red reagent (1:500 dilution) was added to label nuclei of live cells. Cells were imaged using the IncuCyte® live-cell imaging system (Sartorius) with a 10× objective. Images were acquired every 2 hours for up to 72 hours using phase contrast, red (nuclei), and green (GFP-KSHV) channels. Quantification of GFP-positive cells was performed using the integrated IncuCyte analysis software.

### CRISPR/Cas9-mediated knockout of RIG-I

To generate RIG-I–deficient fibroblasts, cells were co-transfected with the human RIG-I CRISPR/Cas9 KO Plasmid (Santa Cruz Biotechnology, sc-400812) and the corresponding homology-directed repair (HDR) plasmid (sc-400812-HDR), using Lipofectamine 3000 (Invitrogen) according to the manufacturer’s protocol. The CRISPR/Cas9 KO plasmid expresses Cas9 and RIG-I targeting guide RNAs, along with a GFP reporter, while the HDR plasmid contains a puromycin resistance cassette and an RFP marker flanked by homology arms. At 48 hours post-transfection, transfected cells were sorted by flow cytometry based on co-expression of GFP and RFP to enrich for successfully edited cells.

### Olink evaluation

RIG-I^WT^ and RIG-I^KO^ fibroblasts (CRISPR-Cas9 edited) were infected with KSHV (MOI = 1) derived from iSLK.219 supernatant. After 72 hours, cell culture supernatants were collected and cleared by centrifugation (500 × g, 5 min), then stored at –80_°C.

Proteins were analyzed using the Olink® Target 96 Inflammation panel (Olink Proteomics AB, Uppsala, Sweden) according to the manufacturer’s instructions. The assay utilizes Proximity Extension Assay (PEA) technology, as detailed by Assarsson et al. (2014) (50) Briefly, antibody probes labeled with oligonucleotides bind to target proteins, and their close proximity enables hybridization of the oligonucleotides. Addition of DNA polymerase triggers a proximity-dependent DNA polymerization event, producing a unique PCR target sequence. These sequences are detected and quantified using the Biomark HD real-time PCR platform (Fluidigm). Data are normalized and quality-controlled, with results expressed as NPX® values—arbitrary units on a log2 scale, where higher values indicate greater protein abundance.

Internal controls used during the PEA process include: two non-human proteins with corresponding antibody probes serve as incubation/immuno controls, an IgG antibody with matching probes functions as the extension control, and a preformed double-stranded amplicon acts as the detection control. These controls are introduced into all samples and external controls. External controls include triplicate negative controls to establish the limit of detection (LOD) and triplicate inter-plate controls (IPCs) containing 92 antibody-probe sets for normalization.

NPX for each sample and assay is calculated using the following steps:1. Ct(analyte) – _Ct(extension control) =dCt(analyte) (to decrease technical variation) 2. dCt(analyte) – _Ct(median IPC)=ddCt(analyte) (to improve inter plate variation) 3. Correction factor(analyte) – _ddCt(analyte) =NPX(analyte) (for more intuitive data). The correction factor is assay- and reagent lot-specific.

Data quality control involves two steps. First, the standard deviation for incubation/immuno controls and detection controls must be below 0.2 for a run to pass. Second, each sample is assessed based on incubation control 2 and the detection control. Samples differing by more than ±0.3 NPX from the plate median for these controls receive a QC warning in the output file. Assay validation data, including detection limits and intra- and inter-assay precision, are available on the manufacturer’s website (www.olink.com). Data analysis was performed using the Olink® Statistical Analysis App (v1.1, 2022-12-23). A Welch’s t-test was applied to compare protein expression between RIG-I^WT^ and RIG-I^KO^ groups for each assay. P-values were adjusted for multiple testing using the Benjamini–Hochberg procedure, with a significance threshold set at adjusted p < 0.05. Proteins with NPX difference > 0.5 and adjusted p < 0.05 were considered differentially expressed. Visualization outputs included hierarchical heatmaps (scaled and clustered), volcano plots, and boxplots.

### RNA-seq analysis

RIG-I^WT^ and RIG-I^KO^ fibroblasts (CRISPR-Cas9–edited) were infected with KSHV for 7 days. Total RNA was extracted using the PureLink™ RNA Mini Kit (Invitrogen), following the manufacturer’s instructions. RNA-seq libraries were prepared using a polyA-enriched, stranded RNA library preparation protocol, and sequenced on an Illumina NovaSeq platform with paired-end 100 bp (PE100) reads. The quality of the raw reads was assessed with FASTQC v0.12.1 and no trimming was deemed necessary. The reads were aligned to the GRCh38 reference genome with STAR v2.7.11b with mean of 93 % of reads uniquely mapped. The raw counts were calculated with FeatureCounts v2.0.6 based on the GRCh38 reference genome (release 110). Differential expression was performed using DESeq2 R package. The top 500 differentially expressed genes (DEGs) from RIG-IWT versus RIG-IKO were kept as an input for enrichment analysis using g:profiler. The top DEGs are defined by rank of adjusted p-value (smallest to largest) x rank of absolute log2 fold changes (largest to smallest) with all genes having an adjusted p-value < 0.05. Results from g:profiler were filtered for enriched terms related to Angiogenesis, Cell proliferation (proliferation or division), Apoptosis (apoptosis or cell death). Related 196 DEGs were selected to draw corresponding heatmap with pheatmap R package, using the z-score of normalized count. In parallel, the full set of top 500 DEGs was analyzed with Metascape (www.metascape.org) to identify broader biological processes affected in RIG-IWT versus RIG-IKO conditions. Metascape output was filtered to display the top 20 significantly enriched GO biological processes among downregulated genes. A network plot was generated in Metascape to visualize functional clusters, where each node represents a GO term and edges reflect shared gene content. Clusters were assigned distinct colors based on semantic similarity to reveal relationships among biological pathways

### Reactivation assays

iSLK.219 cells are a reactivation-competent model harbouring a recombinant latent KSHV genome along with a doxycycline-inducible ORF50 cassette that permits robust entry into the lytic cycle upon stimulation. These cells were transfected with 10 nM siRNA RIG-I or a SiRNA CTL using Lipofectamine RNAiMAX (Thermo Fisher Scientific). At 24 h post-transfection, lytic reactivation was induced with doxycycline (1 µg/mL) for 48 h. Total RNA was extracted using TRIzol reagent (Invitrogen), and KSHV transcript levels were quantified by RT–qPCR. Primary HUVECs and IMR-90 fibroblasts were transfected with 10 nM siRNA CTL or siRNA RIG-I and, 24 h later, infected with KSHV (MOI = 1; derived from iSLK.219 supernatant) by spinoculation (1,500 × g, 1 h, room temperature) in the presence of 8 µg/mL polybrene. After infection, cells were treated with sodium butyrate (NaB, 1 mM) and trichostatin A (TSA, 500 ng/mL) for 24 h to induce histone acetylation and loosen chromatin structure. Unlike iSLK.219 cells, which harbor a doxycycline-inducible ORF50 transgene and support full KSHV reactivation, primary fibroblasts and HUVECs are considered reactivation-incompetent cell types. Following *de novo* infection, these cells rapidly establish latency without detectable ORF50 expression, even upon treatment with chromatin-modifying agents such as TSA or NaB. In the absence of endogenous ORF50 and within an epigenetically restrictive environment, they are unable to initiate a full lytic cascade and instead enter a “pre-lytic” state characterized by partial loosening of latency without productive reactivation(22, 51). For downstream analysis, HUVECs were fixed in 2% paraformaldehyde and stained with ReadyProbes™ F-actin Phalloidin conjugates (Thermo Scientific) according to the manufacturer’s instructions. Images were acquired using the Discover Echo Revolve fluorescence microscope with DAPI (nuclei, blue), FITC (KSHV-GFP, green), and Cy5 (F-actin, red) filter sets. In parallel, total RNA was extracted from iSLK.219 and fibroblasts with TRIzol and analyzed by RT–qPCR using gene-specific primers. Data were normalized to GAPDH.

### Flow Cytometry

Cells were incubated with LIVE/DEAD Fixable Dead Cell Stain (Thermo Fisher Scientific). Samples were analyzed using 4-laser Cytek Aurora flow cytometer (Cytek Biosciences). Data analyses were performed using FlowJo (Ashland, OR). For statistical analysis, repeated-measures two-way ANOVA, followed by Tukey’s multiple comparison post-test were used. The level of significance was set at *p<0.1234, **p<0.0332, ***p<0.0021, ****p<0.0001.

## Acknowledgments

We are extremely grateful to the patient and his family for participating in our research. We thank all the interdisciplinary medical personnel involved in the care of the patient and his family. We thank Shana Lamers and Jasmine Nault from Olink (Thermo Fisher Scientific) for their support.

## Funding

This work was supported by the McGill University Health Centre – Foundation via the SDR Project, through generous support by the Roberts/Hindley family. D.C. Vinh is supported by the Fonds de Recherche du Québec - Santé (FRQS) – Senior clinician-scientist scholar program.

## Author contributions

Conceptualization: DCV, LR Methodology: LR, DCV, VC, SB

Investigation: YS, IU, BM, YL, AW, IA, LRV, AB, SB

Resources: SR, CA, ML, AKW

Funding acquisition: DCV

Supervision: DCV

Writing – original draft: LR, DCV

Writing – review & editing: All authors

## Competing interests

D.C. Vinh has received research support, unrestricted educational grants, consultancy fees, and speaker honoraria from CSL Behring; has received consultancy fees from AstraZeneca, Moderna, and Takeda; and has undertaken clinical trials for Cidara Therapeutics and Moderna. The remaining authors declare no competing financial interests.

## Data and materials availability

All data generated or analyzed during this study are included in this published article and its supplementary information files. Raw data for RNA-seq are available under GEO accession number GSE302318

## Supplementary Materials

Fig. S1. Validation of KSHV latency and reactivation markers in iSLK.219 cells

Fig. S2. RIG-I–dependent IRF response to KSHV in A549 cells

Fig. S3. Validation of CRISPR-Cas9-mediated RIG-I knockout in fibroblasts by Western Blot

Fig. S4. Loss of RIG-I enhances KSHV replication in HMEC

Table S1. Genetic variants previously associated with Kaposi sarcoma

Table S2: Immunophenotyping of peripheral blood mononuclear cells (PBMCs) from patient P1 compared to healthy controls

Table S3. Identified rare, deleterious, recessive genetic variants in P1

## Notes

### Competing Interest Statement

The authors have declared no competing interest.

