## Supplemental Figures/Tables for "Human RIG-I deficiency confers susceptibility to Kaposi Sarcoma via loss of latency control"

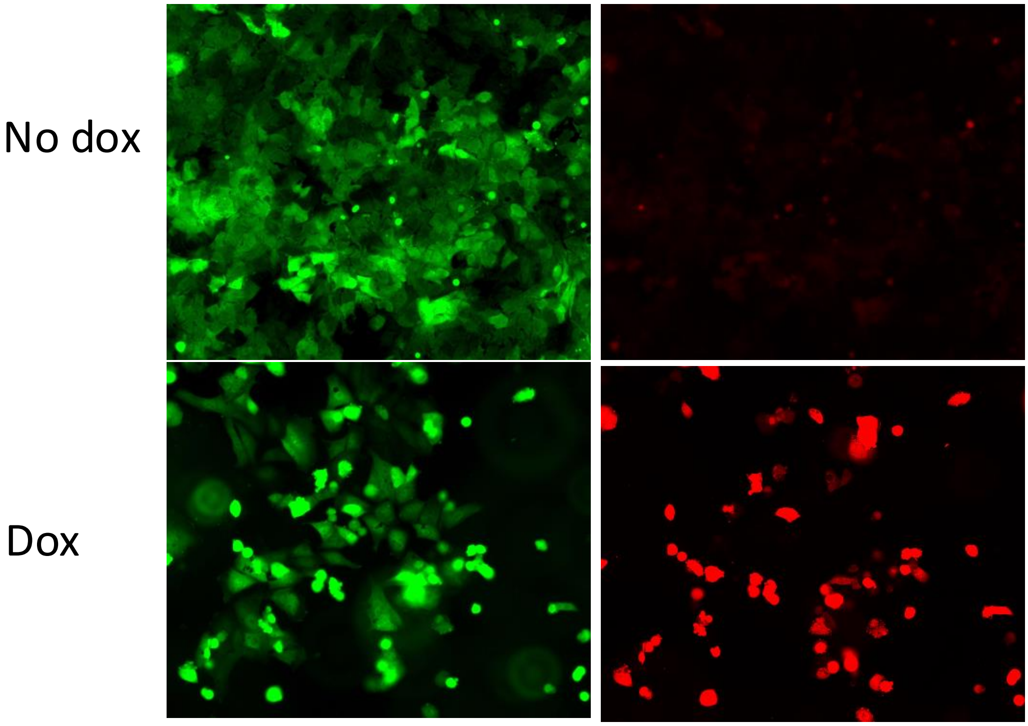

Latent iSLK.219 cells will constitutively express GFP (GFP+),
With Dox : cells enter the lytic cycle (RFA+, GFP+)

**Figure S1: Validation of KSHV latency and reactivation markers in iSLK.219 cells**. In the absence of Dox, iSLK.219 cells maintain latent KSHV infection (GFP+ only). Dox treatment induces the lytic cycle, resulting in dual expression of GFP (latent marker) and RFP (lytic marker, RFP+)

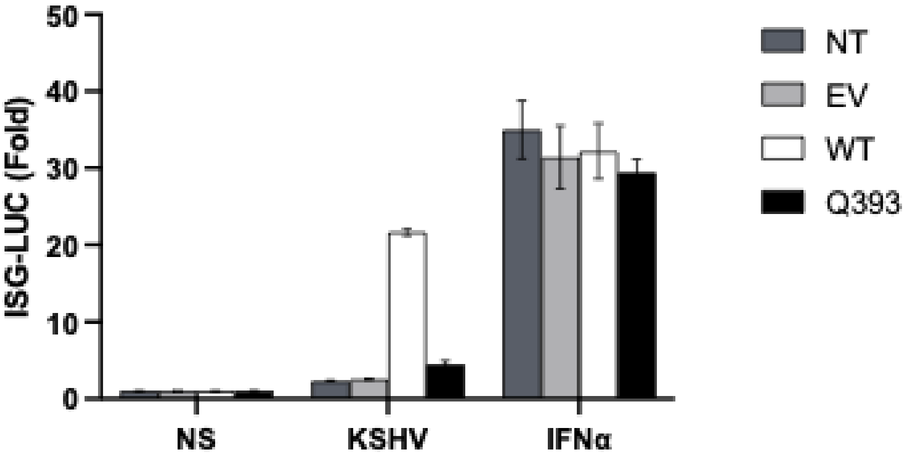

**Figure S2: RIG-I–dependent IRF response to KSHV in A549 cells**. A549-Dual™ KO-RIG-I cells were transfected with either an empty vector (EV), a wild-type RIG-I construct (RIG-I wt) or a RIG-I mutant construct (RIG-I Q393*). Cells were then infected with KSHV or treated with IFNα (positive control) for 24h. Interferon regulatory factor (IRF) pathway activation was assessed by monitoring Lucia luciferase activity. Data are presented as relative luciferase units (RLU). Error bars represent standard deviation of the mean (SD) from triplicate experiments.

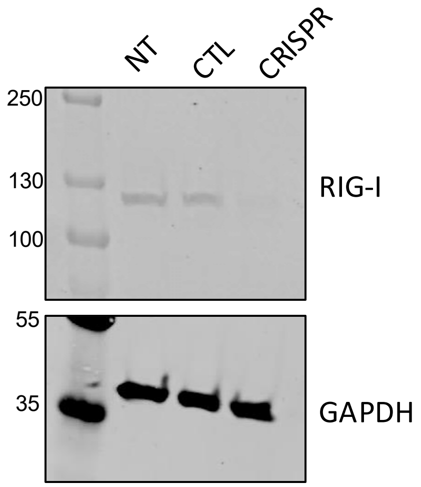

**Figure S3: Validation of CRISPR-Cas9-mediated RIG-I knockout in fibroblasts by Western Blot.** Protein lysates from non-transfected (NT), control plasmid (CTL), and CRISPR-Cas9 RIG-I knockout (CRISPR) fibroblasts were analyzed by Western Blot using an anti-RIG antibody. GAPDH was used as a loading control

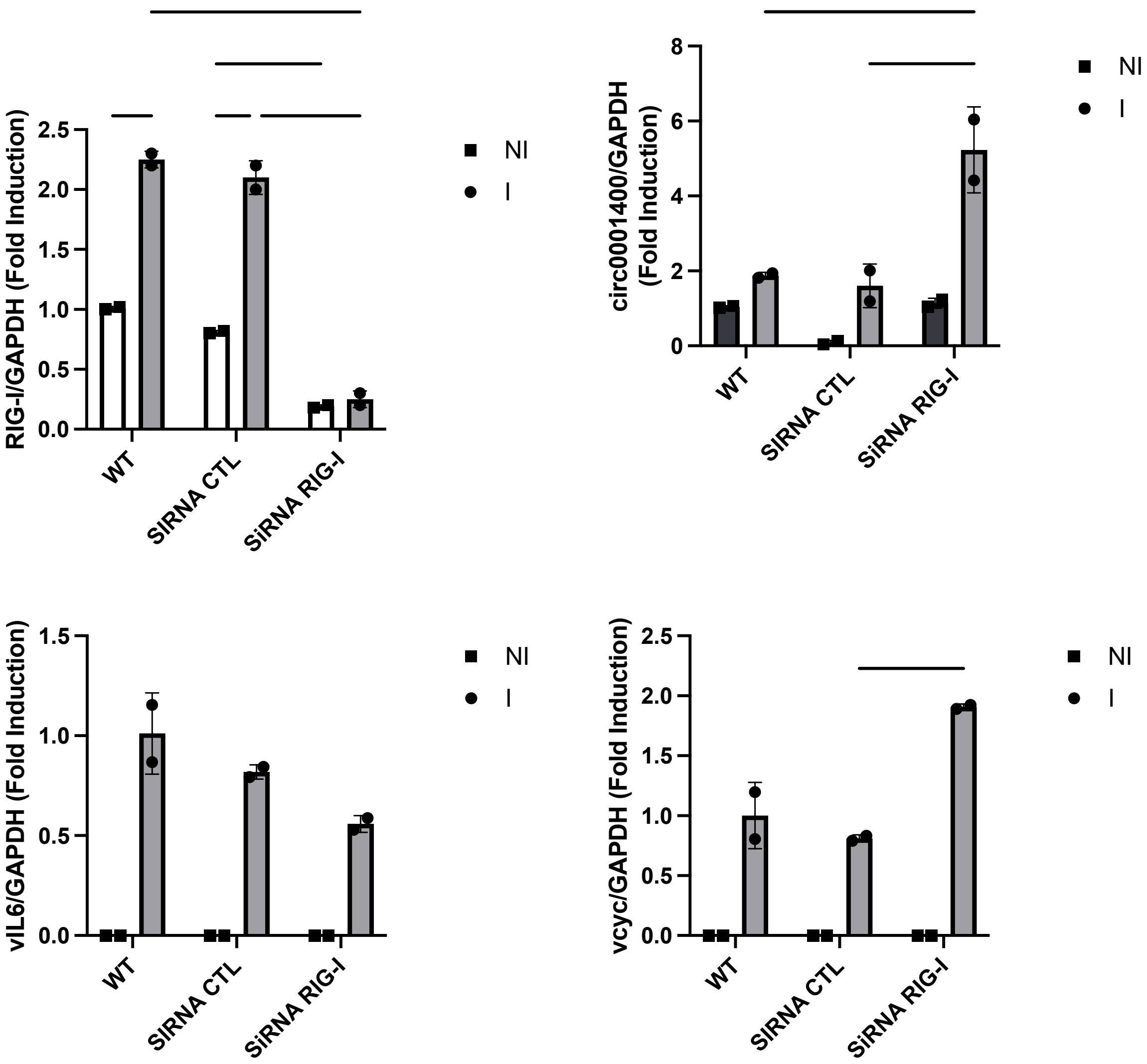

**Figure S4: Loss of RIG-I enhances KSHV replication in HMEC**. HMEC were transfected with either siRNA CTL (control) or siRNA RIG-I and infected with KSHV for 1 week. RNA was extracted, and the expression of KSHV genes (vIL-6, vCyc) and the human-derived circular RNA circ0001400 was analyzed by qPCR. Data are presented as fold induction relative to wild-type non-infected (wt NI) controls, normalized to the housekeeping gene GAPDH. Error bars represent the standard deviation of the mean from triplicate experiments.

### **Table S1:** Genetic variants previously associated with Kaposi sarcoma

| Reference (year) | Age of KS onset / Sex / Ethnicity | Type of KS | Gene / mutation | Defective immunity | Comments |
|---|---|---|---|---|---|
| Camcioglu et al., (2004) | 10 yo/ M/Turkey | Classic Mediterranean | Complete recessive IFNGR1 deficiency / p.C77Y | Impaired IFN-γ response | Disseminated mycobacterial infections. CD4+ T lymphopenia. Disseminated KS; failed IFN-α therapy. Deceased at 12 yo. |
| Picard et al., (2006) | 14 mo/ M/Tunisia | Classic Mediterranean | X-linked recessive WAS / 422del6 | ND | Severe staphylococcal pneumonia. AIHA (treated with steroids). KS occurred 1 month after steroids (for AIHA). Disseminated KS; controlled with taxol. Intracerebral EBV-related lymphoproliferative disorder. Non-T-cell depleted allogeneic HSCT; cured at last follow up. |
| Sahin et al., (2010; case 1); Byun et al., (2010) | 2 yo/ F/Turkey | Classic Mediterranean | Autosomal recessive STIM1 deficiency / 1538-1G>A (leading to loss of expression) | Impaired store-operated Ca2+ entry (T cell deficiency) | AIHA. Disseminated KS. Deceased 4 months after onset. |
| Sahin et al., (2010; case 3); Byun et al., (2013) | 14 yo/ F/Turkey | Classic Mediterranean | Autosomal recessive TNFRSF4 (OX40) deficiency / p.R65C | ↓ OX40 cell surface expression on T cells. ↓ binding of mutant OX40 to its ligand (OX40L). ↓ memory CD4+ T cell function | Previous visceral leishmaniasis (at 9 yo; cured). |
| Aavikko et al., (2015) | 58 yo/ F/Finland | Classic | Heterozygous missense mutation in STAT4 / p.Thr446Ile | No effect on STAT4 phosphorylation (at Y693). ↓ IFN-γ production in naïve Th cells (following PMA + ionomycin) | Indolent: lower extremity, progressing to metastasis. Pulmonary MAC. |
| | 68 yo/ F/Finland | | | | Mentally disabled. Progressive KS lesions. Died of pneumonia at 89 yo |
| | 64 yo/ F/Finland | | | | Died of myocardial infection at 67 yo |
| | 70 yo/ F/Finland | | | | Died of stroke at 73 yo |
| | 86 yo/ F/Finland | | | | Died of myocardial infection at 88 yo |
| Brigida et al., (2017) | 5 yo/ M/Italy | Classic (atypical) | X-MEN disease due to large deletion including MAGT1 (76 Kb deletion, chrX:77,056,603–77,142,993) | ↓ Naïve CD4+ and CD8+ T cells. ↓ memory and switched memory B cells. ↓ surface expression of NKG2D on NK and CD8+ T cells. ↓ NK cells; slight ↓ in NK cytotoxicity to K562. ↓ Ca2+ influx in T cells. | Also had: recurrent respiratory tract infections/bronchiectasis and hypogammaglobulinemia. Atypical feature was presentation with lymphadenopathy (potentially suggesting endemic KS, although classic KS diagnosed based on geographic location and identification of MAGT1 deficiency). Complete remission with peginterferon-α2b and IVIG. |
| Yogev et al., (2023) | Family P1: | Classic | Heterozygous missense mutation in in BPTF / p.I2012T. | Increased expression of LANA, potentially due to interaction between mutant BPTF and LANA. No immunological investigations performed. | |
| | I-2: 81 yo/ F/ Moroccan Jew | | | | I-2: Local disease limited to the lower limbs. Treated with local resection. |
| | II-5: 57 yo / M/ Moroccan Jew | | | | I-5: Advanced classic KS with visceral involvement. Treated with liposomal doxorubicin, vinorelbine, gemcitabine and Alitretinoin cream, local excision, and external beam radiotherapy. |
| | II-10: 44 yo / M/ Moroccan Jew | | | | II-10: Local disease limited to the lower limbs. Treated with local resection. |
| | Family P2 (4 siblings affected; 2 were studied) | | | | |
| | II-3: 49 yo / M/ Moroccan Jew | | | | II-3: Surgical resection. |
| | II-6: 51 yo / M/ Moroccan Jew | | | | II-6: Bilateral leg involvement. Treated with vinblastine and gemcitabine and local external beam radiotherapy. |

**Table S2: Immunophenotyping of peripheral blood mononuclear cells (PBMCs) from patient P1 compared to healthy controls (N=7).**

| Cell Subset | P1 (%) | CTL Median | CTL Range |
|---|---|---|---|
| Monocyte | | | |
| Classical | 29.8 | 71.5 | 66.0-79.6 |
| Intermediate | 2.95 | 5.14 | 0.8-14.1 |
| Lymphocyte | | | |
| **CD3+** | 55.7 | 66.2 | 61.8-78.3 |
| NKT cells | 6.43 | 3.97 | 2.5-10.0 |
| CD8 | 53 | 35.7 | 13.4-46.6 |
| Naïve CD8 | 11.8 | 56.9 | 25.2-62.4 |
| Effector Memory CD8 | 17.9 | 20.6 | 14.1-26.8 |
| Central Memory CD8 | 5.87 | 3.43 | 1.5-9.7 |
| CD4 | 42.6 | 55.9 | 42.1-82.4 |
| Treg | 10.5 | 8.59 | 5.1-37.2 |
| Naïve CD4 | 19.4 | 59.5 | 41.1-68.4 |
| Effector Memory CD4 | 27.9 | 16.3 | 9.7-31.7 |
| Central Memory CD4 | 52.3 | 17.8 | 8.1-32.1 |
| TfH | 12.4 | 7.54 | 1.3-11.5 |
| **CD3-** | 44.3 | 33.8 | 21.6-38.1 |
| CD3-CD19- | 84.3 | 66 | 26.8-81.0 |
| CD56hi | 0.67 | 2.36 | 1.1-7.4 |
| CD56dim=NK | 20.1 | 47.3 | 34.3-80.0 |
| CD56-CD16+ | 5.96 | 10 | 3.0-49.7 |
| CD19+ | 15.1 | 28.9 | 6.9-51.8 |
| Naïve Bcells | 61.9 | 65.5 | 20.6-82.5 |
| IgG+ Memory Bcells | 4.7 | 12.1 | 4.4-16.0 |
| Memory Cells | 20.9 | 19.5 | 9.5-51.1 |
| DC (Lin-HLADR+) | 0.13 | 1.53 | 0.5-3.2 |

Flow cytometry was performed on PBMCs to quantify immune cell subsets in patient P1. Control values represent the median and range (min–max) from seven healthy donors. Values for P1 that fall outside the control range are highlighted in blue.

**Table S3:** Identified rare, deleterious, recessive genetic variants in P1

| Genomic Position (hg38) | Reference/Alternate Alleles | Allele Frequency (Patient) | Gene | gnomAD Allele Frequency | CADD_PHRED |
|---|---|---|---|---|---|
| chr3:195507323 | T/TGACCTGTGGATGATGAGGAAGGGCTGGTGACATGAAGAGGGGTGACGTGACCTGTAGATACTGAGGAAGTGCTGGTGACAGGAAGAGGGGTGGG | 1 | MUC4 | 0 | None |
| chrX:49454149 | TAAAAAAAAAAAAAAAAAAAAAA/T | 1 | PAGE1 | 0.00096 | None |
| chr9:32487978 | G/A | 1 | DDX58 | 0 | 40 |
| chr5:93722109 | C/T | 1 | KIAA0825 | 0 | 33 |
| chr3:143212596 | A/C | 1 | SLC9A9 | 0 | 31 |
| chr2:219528715 | C/T | 1 | RNF25 | 0 | 29 |
| chr12:113724871 | G/A | 1 | TPCN1 | 0.000074 | 28.2 |
| chrX:19020970 | C/T | 1 | ADGRG2 | 0.00005 | 28.1 |
| chr5:154181823 | G/C | 1 | LARP1 | 0.00012 | 27.3 |
| chr2:48062009 | T/C | 1 | FBXO11,IGKC | 0.0000081 | 26.2 |
| chr17:76430187 | G/A | 1 | DNAH17 | 0.000092 | 26.2 |
| chr16:30392742 | C/G | 1 | MYLPF,ZNF48,SEPTIN1 | 0.000024 | 25.5 |
| chr11:68548130 | G/A | 1 | CPT1A | 0.000032 | 24.8 |
| chr12:43821148 | C/T | 1 | ADAMTS20 | 0.000053 | 24.4 |
| chr2:231949791 | G/C | 1 | PSMD1 | 0 | 24.2 |
| chr22:41077844 | C/T | 1 | MCHR1 | 0.000076 | 23.7 |
| chr2:137814296 | T/C | 1 | THSD7B,IGKC | 0.00022 | 23.5 |
| chr13:79940840 | G/T | 1 | RBM26 | 0.000016 | 23.3 |
| chrX:106843700 | C/G | 1 | FRMPD3 | 0 | 23 |
| chr13:21746644 | C/CGCTTTCCTCTCCGTCCAGCGCTCACTGCAGCCGGGCCG | 0.78 | MRPL57,SKA3 | 0.000059 | 22.7 |
| chr13:21746644 | C/CGCTTTCCTCTCCGTCCAGCGCTCACTGCAGCCGGGCCGTCTCGCAG | 0.89 | MRPL57,SKA3 | 0.00013 | 22.6 |
| chr15:59516890 | G/C | 1 | MYO1E | 0.000085 | 22.4 |
| chr6:32551942 | T/C | 1 | HLA-DRB1 | 0.000027 | 21.9 |
| chrX:57020758 | C/T | 1 | SPIN3 | 0 | 19 |
| chr1:119530498 | C/A | 1 | TBX15 | 0.00013 | 18.06 |
| chr19:57325555 | TTGGCTCAGCAGCCTCCACTTC/T | 1 | PEG3 | 0.00052 | 17.95 |
| chr3:57542866 | G/A | 1 | PDE12 | 0.000011 | 17.34 |
| chr1:200708987 | G/A | 1 | CAMSAP2 | 0.000036 | 17.24 |
| chr12:123032058 | TCTC/T | 1 | KNTC1 | 0.00096 | 15.01 |
| chr17:8661598 | G/A | 1 | SPDYE4 | 0.00006 | 14.48 |
| chrX:152770776 | A/G | 1 | BGN | 0 | 14.44 |
| chrX:108695343 | A/G | 1 | GUCY2F | 0.000011 | 14.33 |
| chr22:41726113 | T/C | 1 | ZC3H7B | 0.000004 | 13.65 |
| chr17:8925629 | C/A | 1 | NTN1 | 0.000096 | 13.24 |
| chr12:113796419 | T/G | 1 | PLBD2 | 0.000014 | 12.41 |
| chr10:134121262 | G/T | 0.74 | STK32C | 0 | 11.91 |
| chr19:35832255 | G/A | 0.4 | CD22 | 0.000018 | 17.22 |
| chr19:35832331 | C/A | 0.47 | CD22 | 0.000046 | 22.6 |
| chr3:195508387 | G/A | 0.22 | MUC4 | 0 | 15.4 |
| chr3:195507323 | T/TGACCTGTGGATGATGAGGAAGGGCTGGTGACATGAAGAGGGGTGACGTGACCTGTAGATACTGAGGAAGTGCTGGTGACAGGAAGAGGGGTGGG | 1 | MUC4 | 0 | None |
| chr2:152483562 | G/A | 0.46 | NEB | 0 | 23.8 |
| chr2:152490404 | T/C | 0.48 | NEB | 0.00039 | 19.39 |
| chr7:98256631 | G/A | 0.51 | NPTX2 | 0 | 27.8 |
| chr7:98247072 | G/T | 0.52 | NPTX2 | 0.0004 | 20.4 |
| chr1:33236735 | C/A | 0.44 | NHSL3 | 0 | 11.53 |
| chr1:33235862 | C/T | 0.48 | NHSL3 | 0.000069 | 27.1 |
